# Down the Penrose stairs: How selection for fewer recombination hotspots maintains their existence

**DOI:** 10.1101/2022.09.27.509707

**Authors:** Zachary Baker, Molly Przeworski, Guy Sella

## Abstract

In many species, meiotic recombination events tend to occur in narrow intervals of the genome, known as hotspots. In humans and mice, double strand break (DSB) hotspot locations are determined by the DNA-binding specificity of the zinc finger array of the PRDM9 protein, which is rapidly evolving at residues in contact with DNA. Previous models explained this rapid evolution in terms of the need to restore PRDM9 binding sites lost to gene conversion over time, under the assumption that more PRDM9 binding always leads to more DSBs. In recent experimental work, however, it has become apparent that PRDM9 binding on both homologs facilitates DSB repair, and moreover that, in the absence of enough symmetric binding, meiosis no longer progresses reliably. We therefore consider the possibility that the benefit of PRDM9 stems from its role in coupling DSB formation and efficient repair. To this end, we model the evolution of PRDM9 from first principles: from its binding dynamics to the population processes that govern the evolution of the zinc finger array and its binding sites in the genome. As we show, the loss of a small number of strong binding sites leads to the use of a greater number of weaker ones, resulting in a sharp reduction in symmetric binding and favoring new PRDM9 alleles that restore the use of a smaller set of strong binding sites. This decrease in PRDM9 binding symmetry and in its ability to promote DSB repair drive the rapid zinc finger turnover. These results imply that the advantage of new PRDM9 alleles is in *limiting* the number of binding sites used effectively, rather than in increasing net PRDM9 binding, as previously believed. By extension, our model suggests that the evolutionary advantage of hotspots may have been to increase the efficiency of DSB repair and/or homolog pairing.

## Introduction

Meiotic recombination is initiated by the deliberate infliction of double strand breaks (DSBs) across the genome and resolved with their subsequent repair off homologous chromosomes. In many species, including all plants, fungi and vertebrates investigated to date, the vast majority of meiotic recombination events localize to narrow intervals of the genome known as *hotspots* (reviewed in Tock and Henderson 2018). Why recombination is concentrated in hotspots in some taxa, as opposed to being more uniformly spread across the genome, as in others, such as *Drosophila* (Manzano-Winkler et al. 2013; Smukowski et al. 2015) or *C. elegans* (Kaur et al. 2014; Bernstein et al. 2016), remains unclear. Moreover, among species with hotspots, there exists a great deal of variation in how hotspots are localized, and it is largely unknown why these different mechanisms have been adopted in different lineages.

In humans and mice, and likely many other metazoan species, the primary determinant of hotspot locations is PRDM9 and the DNA-binding specificity of its Cys-2-His-2 Zinc Finger (ZF) array (Myers et al. 2010; Baudat et al. 2010; Parvanov et al. 2010; Baker et al. 2017). Early in meiosis, PRDM9 binds to thousands of binding sites along the genome (Grey et al. 2017; Parvanov et al. 2017), where its PR/SET domain catalyzes the formation of H3K4me3 and H3K36me3 marks on nearby histones (Hayashi et al. 2005; Grey et al. 2011; Wu et al. 2013; Eram et al. 2014; Powers et al. 2016). In conjunction with protein interactions mediated by the KRAB domain of PRDM9, these marks play a role in bringing PRDM9-bound sites to the chromosomal axis, where the SPO11 protein catalyzes the formation of DSBs at a small subset (Imai et al. 2017; Parvanov et al. 2017; Diagouraga et al. 2018; Thibault-Sennett et al. 2018; Bhattacharyya et al. 2018).

While PRDM9-mediated recombination is concentrated away from promoter-like features of the genome such as transcriptional start sites or CpG islands (Brick et al. 2012; Baker et al. 2017), in many species lacking PRDM9 genes, such as canids, birds, fungi and plants, recombination is concentrated near such features (Auton et al. 2013; Singal et al. 2015; Lam and Keeney 2015; Yelina et al. 2015). The same is observed in PRDM9-/- mice (Brick et al. 2012) and, more tentatively, in a human female lacking both copies of PRDM9 (Narasimhan et al. 2016). This observation suggests that in species that habitually use PRDM9, this mechanism masks a “default” mechanism for recombination that resembles the one employed by species lacking PRDM9.

Intriguingly, the use of promoter-like features is associated with conserved recombination landscapes over tens of millions of years (Singal et al. 2015; Lam and Keeney 2015; Liu et al. 2019), whereas in species that use PRDM9 to direct recombination, the recombination landscape is rapidly-evolving. Consequently, closely related species, such as humans and chimpanzees, or even different subspecies, use sets of hotspots with no more overlap than expected by chance (Ptak et al 2004; Winckler et al 2005; Auton et al. 2012; Brick et al. 2012; Stevison et al. 2016). This rapid evolution is thought to result from both the evolutionary erosion of PRDM9 binding sites due to gene conversion (GC) and repeated changes to the DNA-binding specificity of PRDM9’s ZF-array.

More specifically, when individuals are heterozygous for “hot” and “cold” alleles, defined as those more or less likely to experience a DSB during meiosis, GC leads to the under-transmission of hot alleles in a manner mathematically analogous to purifying selection against them (Nagalaki and Petes 1982; Jeffreys and Neumann 2002). This phenomenon can be observed in the over-transmission of polymorphisms disrupting PRDM9 binding motifs within hotspots (Berg et al. 2011; Cole et al. 2014). In the absence of any countervailing selective pressures, the under-transmission of hot alleles should lead to their rapid loss from the population over time (Nicolas et al. 1989; Boulton et al. 1997; Pineda-Krch and Redfield 2005; Coop and Myers 2007; Peters 2008); this too is observed, in the unusually rapid erosion of PRDM9 binding sites in primate and rodent lineages (Myers *et al*., 2010; Baker *et al*., 2015; Smagulova *et al*., 2016; Spence and Song 2019).

Despite the losses of individual hotspots, hotspots persist in the genome, presumably as changes in the DNA-binding specificity of PRDM9 give rise to new sets of hotspots. Variation in hotspot usage has been associated with different PRDM9 alleles (i.e., with different amino acids at the DNA-binding residues of their ZFs) in mice, humans and cattle (Myers et al. 2010; Baudat et al. 2010; Parvanov et al. 2010; Berg et al. 2010; Kong et al. 2010; Berg et al. 2011; Fledel-Alon et al. 2011; Hinch et al. 2012; Sandor et al. 2012; Ma et al. 2015), and the experimental introduction of a PRDM9 allele with a novel ZF-array into mice reprograms the locations of hotspots (Davies et al. 2016). Moreover, strong signatures of positive selection on the DNA-binding residues of ZFs from PRDM9 genes suggests a rapid turnover of PRDM9 binding specificity driven by recurrent selection for novel binding specificities (Oliver et al. 2009; Thomas et al. 2009; Myers et al. 2010; Buard et al. 2014; Baker et al., 2017).

To make sense of these observations, a number of theoretical models have been developed that describe the co-evolution of PRDM9 and its binding sites (Úbeda and Wilkins 2011; Latrille et al. 2017). In these models, GC acts to remove PRDM9 binding sites over time, leading to a proportional reduction in the amount of PRDM9 bound and consequently of DSBs or crossovers (COs). Under the assumption that having too few DSBs or COs leads to a decrease in fitness, potentially as a consequence of a requirement for one CO per chromosome for proper disjunction, the loss of binding sites and resulting recombination events is deleterious. These models thus give rise to a *Red Queen* dynamic, in which younger PRDM9 alleles are favored over older ones because their binding sites have experienced less erosion due to GC.

These models demonstrate how the use of hotspots can persist despite GC against hotter alleles, by coupling the evolution of PRDM9 to the loss of its binding sites. However, arguing that new PRDM9 alleles are favored because they restore PRDM9 binding seems at odds with the nature of hotspots, namely the use of only a small proportion of the genome for recombination. Taken to the extreme, if selection simply favors PRDM9 alleles with a greater number of binding sites, an optimal PRDM9 allele will be one that can bind anywhere in the genome, at which point the heats of individual binding sites would be so diluted as to no longer be considered hotspots.

Moreover, multiple lines of evidence suggest that the loss of PRDM9 binding sites in the genome does not in fact imperil the initiation of recombination events, but merely alters their locations. In particular, the number of DSBs formed per meiosis is tightly regulated independently of PRDM9 (Kauppi et al. 2013; Yamada et al. 2017); similar numbers of DSBs form in PRDM9-/- mice (Hayashi et al. 2005; Brick et al. 2012; Mihola et al. 2019); and PRDM9-bound sites appear to compete for DSBs (Diagouraga et al. 2018). Furthermore, PRDM9 binding sites themselves are known to compete for available PRDM9 molecules, such that the loss of binding sites may only weakly affect the number bound (Billings et al. 2013, Baker et al. 2014).

In addition to serving to localize DSBs, PRDM9 is now appreciated to play a role in DSB repair, mediated in part by its symmetric recruitment to the same loci on homologous chromosomes during meiosis (Smagulova et al. 2016; Davies et al. 2016; Gregorova et al. 2018; Hinch et al. 2019; Li et al. 2019; Huang et al. 2020; Mahgroub et al. 2020; Wells et al. 2020; Gergelits et al. 2021; Davies et al. 2021). This secondary function was first suggested by studies of hybrid sterility in mice, in which particular crosses result in the production of sterile F1 males (reviewed in Forejt et al. 2021). In sterile hybrids, the PRDM9 allele of each parental subspecies preferentially binds to the non-parental background, as a consequence of the strongest PRDM9 binding sites in each parental subspecies having eroded in a lineage-specific manner. The resulting asymmetric binding leads to meiotic arrest in pachytene associated with widespread delays in DSB repair, asynaptic chromosomes, and defects in homolog pairing (Baker et al. 2015; Smagulova et al. 2016; Davies et al. 2016). Restoring PRDM9 binding symmetry, either through the introduction of a novel PRDM9 allele with binding sites not yet eroded in either parental subspecies (Davies et al. 2016; Davies et al. 2021), or by partially restoring the homozygosity of binding sites on the chromosomes most prone to asynapsis (Gregorova et al. 2018), is sufficient to rescue the fertility of hybrids. These results thus indicate that fertility can be restored when a small number of DSBs are made at sites symmetrically bound by PRDM9 (Gregorova et al. 2018) on each chromosome.

The specific importance of PRDM9 binding symmetrically likely stems from the fact such sites can be repaired more rapidly, and that early recombining sites are preferentially used for COs and homolog pairing. In particular, genome-wide analyses of DSB outcomes at PRDM9 binding sites provide direct evidence that DSBs at asymmetrically-bound hotspots are more likely to experience delays in their repair (Hinch et al. 2019; Li et al. 2019), and further reveal that, possibly because of this delay, they are less likely to result in COs (Hinch et al. 2019), and more likely to be resolved using the sister chromatid as a template instead of the homologous chromosome (Li et al. 2019; Gergelits et al. 2021). Recent work has further implicated the gene *ZCWPW1* as a key player in mediating the efficient repair of sites bound symmetrically by PRDM9. Namely, ZCWPW1 binds PRDM9-mediated histone marks on the unbroken homolog, leading to their preferential use as templates for DSB repair, independent of their preferential use as sites of DSB initiation (Huang et al. 2020, Mahgroub et al. 2020, Wells et al. 2020). Taken together, these findings indicate that PRDM9 binding symmetry increases the efficiency of DSB repair, and possibly, of homolog pairing.

These observations raise the possibility that selection driving the evolution of the PRDM9 ZF stems from its role in DSB repair, rather than DSB initiation, as previously envisioned. The *Red Queen* theory of recombination hotspots suggests that the erosion of PRDM9 binding sites should lead to a measurable decrease in fitness, which can be restored by the introduction of a novel PRDM9 allele. The asymmetric binding and associated reduced fertility observed in hybrid mice is the only known evidence consistent with this phenomenon. It occurs as the consequence of the lineage-specific erosion in each parental lineage leading to widespread heterozygosity at PRDM9 binding sites, an extreme scenario that is unlikely to occur within populations, in part because gene conversion will act to rapidly remove heterozygosity at PRDM9 binding sites. Instead, one might envisage that PRDM9 binding asymmetry arises not from heterozygosity in its underlying binding sites, but as a consequence of PRDM9’s competitive binding: namely, that the erosion of strong binding sites shifts binding toward many more weaker ones, and in so doing reduces the chances of symmetric binding and of the successful completion of meiosis.

On that basis, it seems likely that selection on PRDM9 is mediated by a requirement for symmetric binding. Based on this premise, we model the co-evolution of PRDM9 and its binding sites from first principles, assuming, as seems more realistic, that: (i) the number of DSBs is regulated independently of PRDM9, (ii) PRDM9 binding is competitive between sites, and (iii) fitness is a function of the probability that the smallest chromosome experiences some minimal number of DSBs at sites symmetrically bound by PRDM9.

## The Model

### Overview

We model the co-evolution of PRDM9 and its binding sites in a panmictic population of constant size. An individual is described in terms of their two PRDM9 alleles. Each PRDM9 allele recognizes a distinct set of binding sites, which may have distinct binding affinities or “heats” (i.e., different probabilities of being bound). The model is composed of four main parts (**Fig. 1**). The first describes PRDM9 binding during meiosis in terms of Michaelis-Menten kinetics, in which sites compete for binding their cognate PRDM9 proteins. We use this model to approximate the probability that any given site on the four chromatids (i.e., the four copies of a chromosome present during meiosis) is bound by PRDM9 (**Fig 1A**). We then approximate the probabilities of any given site experiencing a DSB, and of any given locus experiencing a DSB while being symmetrically bound by PRDM9, i.e., of a *symmetric DSB* (**Fig. 1B**).

**Figure 1.**
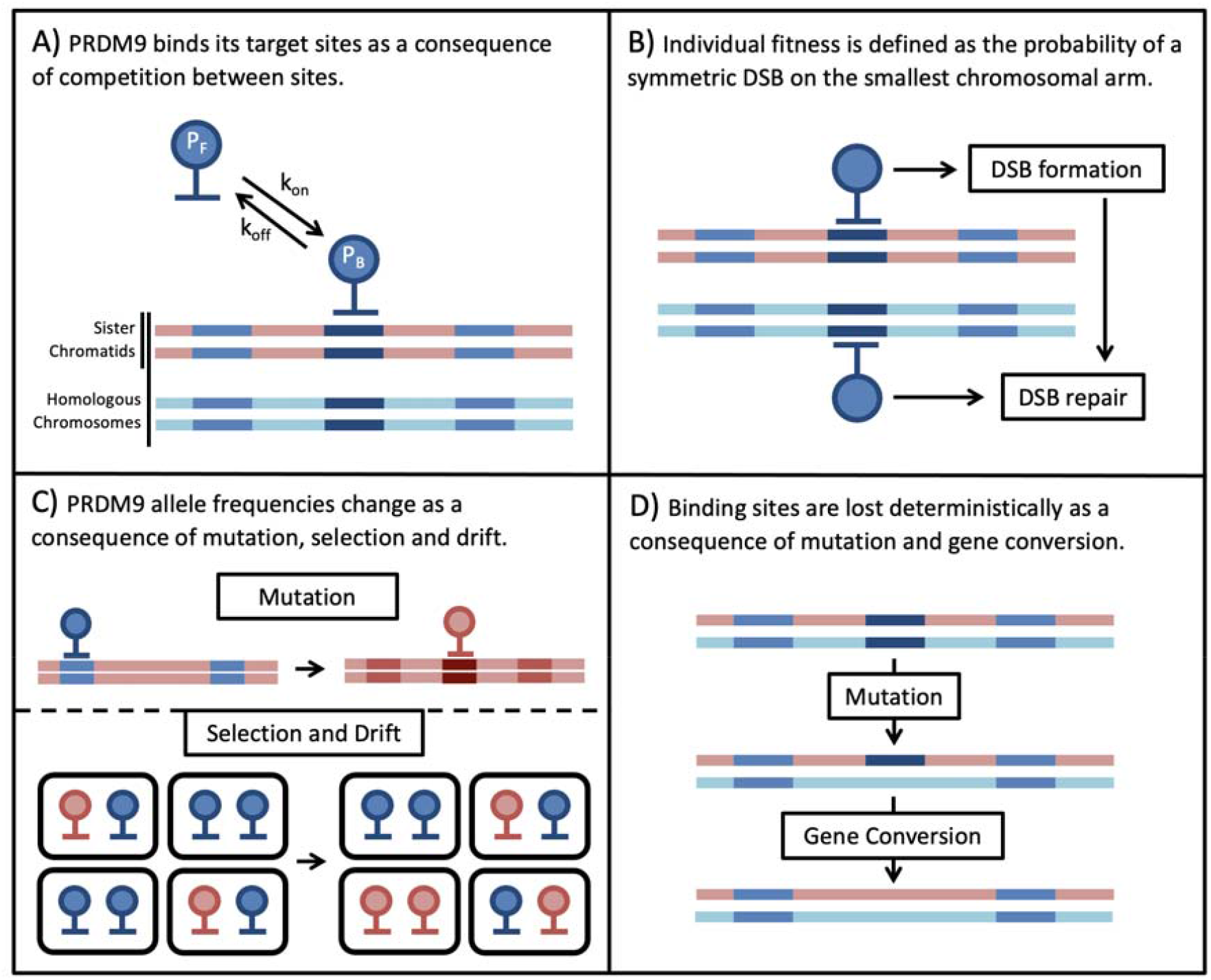
Overview of the Mode.

Next, we model the population processes. We use Wright-Fisher sampling with viability selection in a panmictic diploid population of size to describe the change in PRDM9 allele frequencies in the population (**Fig. 1C**). New PRDM9 alleles arise from mutation with probability per individual meiosis and establish a new set of binding sites. Selection on a PRDM9 allele arises from its impact on fitness, and more specifically, the need for at least one symmetric DSB on per chromosome per meiosis. For simplicity, we consider only the smallest chromosome because fitness defects arising from asymmetry in PRDM9 binding should affect the smallest chromosomes first (assuming, as seems plausible, that they have fewer sites at which a symmetric DSB might be made). Also for simplicity, we assume that individual fitness is equal to the probability of a symmetric DSB in a single meiosis. In reality, we expect fitness to depend on the proportion of an individual’s gametes that have a sufficient number of symmetric DSBs, but this fitness should be a monotonically-increasing function of the probability per meiosis, and therefore we expect the qualitative conclusions to be unchanged. Lastly, we use a deterministic approximation to describe the erosion of binding sites over time (**Fig. 1D**). Mutations that destroy the ability of a site to bind to its cognate PRDM9 are introduced with probability *µ* per individual meiosis and can fix in the population due to biased gene conversion. The fixation probability depends on rates at which the binding site experiences a DSB and of GC. Below, we describe each part of the model in turn.

### Molecular dynamics of PRDM9 binding

We model the binding of PRDM9 as a consequence of Michaelis-Menten kinetics, wherein binding sites compete for association with a fixed number of PRDM9 molecules. In **Supplementary Section S1**, we solve these dynamics at equilibrium, i.e., when the rate at which PRDM9 dissociates from sites with any given binding affinity (“heat”) is equal to the rate at which PRDM9 binds those sites. Specifically, we show that the *heat* of a site, i.e., the probability that a site with binding affinity is bound by PRDM, can be expressed as

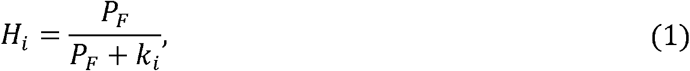

where *k*_*i*_ is the site’s dissociation constant and *P*_*F*_ is the number of free PRDM9 molecules. In turn, we show that the expected number of free PRDM9 molecules satisfies the equation

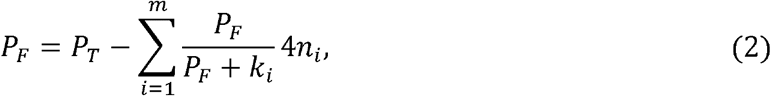

where *P*_*T*_ is the total number of PRDM9 molecules, *m* is the number of binding affinities, and *n*_*i*_ is the number of sites with affinity *i*, with *i* = 1 … *m*. When we consider models with one or two types of binding sites, this equation becomes a quadratic or cubic in *P*_*F*_, respectively, which we solve analytically (see **Supplementary Section S1**). In the general case, this equation becomes a polynomial of degree *m* + 1, which can be solved numerically.

The total number of PRDM9 proteins of a given type depends on whether an individual is homozygous or heterozygous for the corresponding PRDM9 allele. We assume that a fixed number of PRDM9 molecules are produced per allele. Consequently, the corresponding PRDM9 binding sites will always be more likely to be bound in individuals homozygous for the cognate PRDM9 allele relative to heterozygotes. Moreover, the ratio of heats for sites with different binding affinities will differ between genotypes.

### Molecular dynamics of DSBs and symmetric DSBs

Next, we approximate the probability that a PRDM9-bound site experiences a DSB in a given meiosis. We assume that the total number of DSBs (*D*) is constant and that all PRDM9 bound sites are equally likely to experience a DSB. Thus, if the total number of PRDM9-bound sites in an individual with PRDM9 alleles *j* and *k* is 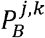 then the probability that a bound site experiences a DSB is

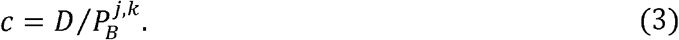

This approximation assumes that conditional on being bound, PRDM9 molecules arising from different alleles are equally likely to recruit the DSB machinery, but allows for some degree of dominance as a consequence of differences in the numbers of sites bound by each allelic variant genome-wide.

In **Supplementary Section S2**, we show that the probability that a site with binding affinity *i* experiences a symmetric DSB, i.e., that at least one of the chromatids is bound by PRDM9 and experiences a DSB and that at least one sister chromatid of each homolog is bound by PRDM9, is.

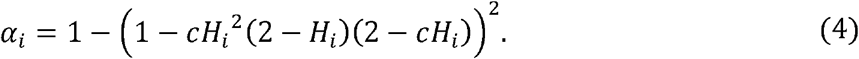

For brevity, we omitted indexes corresponding to the genotype at the PRDM9 locus, which affect *α*_*i*_ through *c* (Eq.3) and *H*_*i*_ through *P*_*T*_ (Eq. 2).

### Evolutionary dynamics of PRDM9 alleles

We model individual fitness as the probability per meiosis that at least one symmetric DSB occurs on the smallest chromosome, which comprises a proportion *r* of the genome. We make the simplifying assumption that the proportion of binding sites of any affinity found on that chromosome is also *r*, and remains fixed over time; the number of binding sites with binding affinity *i* is therefore *rn*_*i*_. We also assume that the probability of symmetric DSBs at different binding sites are independent (i.e., ignoring interference between them). Under these assumptions, we approximate the fitness of a homozygote for PRDM9 allele *j* as one minus the probability that none of the binding sites on the smallest chromosome experience a symmetric DSB, i.e.,

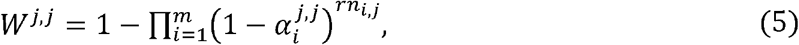

where the product is taken over the *m* possible binding affinities, and *n*_*i,j*_ is the number of binding sites with affinity *i* associated with PRDM9 allele *j*. Similarly, we approximate the fitness of a heterozygous individual for PRDM9 with alleles *j* and *k* ≠ *j*, as

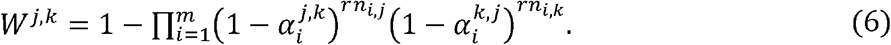

The marginal fitness associated with a PRDM9 allele is calculated as a weighted average over genotypic frequencies, in which homozygotes for the focal allele, *j*, are counted twice to account for the two copies of the allele, i.e.,

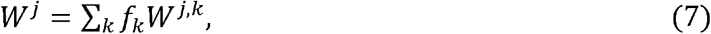

where *f*_*k*_ is the frequency of PRDM9 allele *k*. The expected frequencies of allele *j* after viability selection is then given by

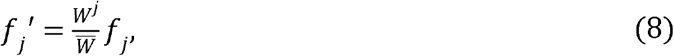

where 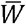 denotes the population’s mean fitness. These expectations are used in Wright-Fisher sampling across generations.

### Evolutionary dynamics of PRDM9 binding sites

We model the loss of PRDM9 binding sites due to biased gene conversion deterministically. Cold alleles at a binding site arise in the population at a rate of *2N*μ per generation, where *N* is the population size and μ is the mutation rate from a hot to a cold allele per gamete per generation. Biased gene conversion acts analogously to selection against hot alleles with a selection coefficient equal to the rate of gene conversion fully spanning that binding site, *g* (Nagylaki & Petes 1982). Consequently, a newly arisen cold allele (which is never bound by PRDM9) will eventually fix in the population with probability 2*g*. In turn, we model the probability 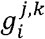 that a binding site with affinity *i* and recognized by PRDM9 allele *j* experiences gene conversion during a given meiosis in an individual with PRDM9 alleles *j* and *k* as the product of the probabilities that it is bound by PRDM9 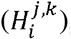, experiences a DSB conditional on being bound (*c*^*j,k*^), and that the resulting gene conversion spans the binding site (*B*), i.e.,.

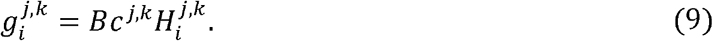

The expected population rate of gene conversion at this binding site is the average over individual rates, weighted by genotype frequencies (after viability selection), i.e.,

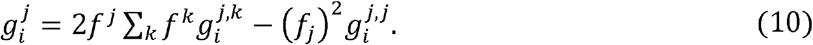

We model the fixations of cold alleles as if they occur instantaneously after they arise as mutations, with probabilities that derive from the rate of gene conversion at that time. Thus, we approximate the reduction in the number of binding sites with affinity *i* for PRDM9 allele *j* in a given generation by

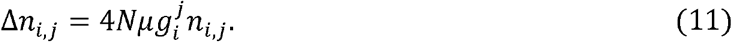

We note that because we assume that the number of DSBs and thus of gene conversion events per generation is constant, the total rate at which binding sites are lost, across all heats and PRDM9 alleles, is constant, and equals

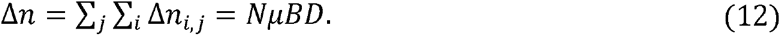

### Simulations

Simulations of the model were implemented using a custom R script (available at https://github.com/sellalab). The simulation keeps track of the frequencies of PRDM9 alleles and of the numbers of binding sites of each considered heat for each allele. Each generation, the simulation calculates the marginal fitness of each PRDM9 allele (Eq. 7) and the number of sites of each heat lost due to gene conversion (Eq. 11). The frequencies of PRDM9 alleles for the succeeding generation are generated by Wright-Fisher (multinomial) sampling with viability selection (Eq. 8). The number of new PRDM9 alleles generated by mutation each generation is sampled from a binomial distribution, where the probability of success is the mutation rate at PRDM9 *v*, and the number of trials is 2*N*. These mutations are randomly chosen to replace pre-existing PRDM9 alleles (whose frequencies are correspondingly decreased by 1/2*N* per mutation). In every generation, the program records various quantities discussed below, such as the population average fitness or heterozygosity at the PRDM9 locus.

### Estimated parameters

Except where stated otherwise, the parameters used in simulations are given in **Table 1**. The mutation rate from hot to cold alleles is taken to be 1.25 × 10^−7^ per gamete per generation, to reflect a per base pair rate of 1.25 × 10^−8^, as estimated in humans (Kong et al. 2012), and ~10 non-degenerate base pairs for PRDM9 binding per binding site, roughly consistent with observed binding motifs in mice and humans (Baudat et al. 2010; Myers et al. 2010). The mutation rate of new PRDM9 alleles is taken to be 10^−5^ per gamete per generation, based on observations in humans (Jeffreys et al., 2013). The number of DSBs per meiosis is set to 300, in light of estimates in both humans and mice (Baudat et al., 2007). The number of PRDM9 molecules expressed during meiosis is set to 5,000, roughly consistent with cytological observations of thousands of PRDM9 foci in the nuclei of meiotic cells from mice (Parvanov et al. 2017; Grey et al. 2017), under an assumption that most PRDM9 molecules are bound. We set the probability that a gene conversion tract spans the PRDM9 binding motif to 0.7, based on estimates from mice (Li et al. 2019). Lastly, we set the proportion of the genome corresponding to the smallest chromosome to be 1/40, roughly equivalent to that of the smallest chromosome in mice.

**Table 1.** Parameters and variables of the model.

| *Parameter* | *Description* | *Value* |
| --- | --- | --- |
| $N$ | Diploid population size | variable |
| $\mu$ | Mutation rate from hot to cold alleles at PRDM9 binding sites per generation | $1.25 \times 10^{-7}$ |
| $\nu$ | Mutation rate at the PRDM9 locus per generation | $10^{-5}$ |
| $D$ | Number of DSBs initiated per meiosis | 300 |
| $P_T$ | Total number of PRDM9 molecules expressed per meiosis | 5,000 |
| $k_i$ | Dissociation constant of PRDM9 binding sites with affinity $i$ | variable |
| $B$ | Probability that a conversion tract spans the PRDM9 binding motif | 0.7 |
| $r$ | Proportion of the genome corresponding to the smallest chromosome | 1/40 |
| *Variable* | *Description* |  |
| $n_i^j$ | Number of PRDM9 binding sites with affinity $i$, recognized by PRDM9 allele $j$, across the genome | |
| $f^j$ | The allele frequency of PRDM9 allele $j$ | |
| $P_F^{j,k}$ | Number of free cognate PRDM9 molecules in an individual with PRDM9 alleles $j$ and $k$ per meiosis | |
| $P_B^{j,k}$ | Total number of bound cognate PRDM9 molecules in an individual with PRDM9 alleles $j$ and $k$ | |
| $H_i^{j,k}$ | Probability that a given site with binding affinity $i$, recognized by PRDM9 allele $j$, is bound by PRDM9 in an individual with PRDM9 alleles $j$ and $k$. | |
| $c^{j,k}$ | Probability that any PRDM9-bound site experiences a DSB in an individual with PRDM9 alleles $j$ and $k$. | |
| $\alpha_i^{j,k}$ | Probability of a PRDM9 binding site with affinity $i$, recognized by PRDM9 allele $j$, being symmetrically bound by PRDM9 and experiencing a DSB in an individual with PRDM9 alleles $j$ and $k$. | |
| $W^{j,k}$ | The fitness of an individual with PRDM9 alleles $j$ and $k$ | |
| $W^j$ | The marginal fitness of PRDM9 allele $j$ across possible genotypes | |
| $g_i^{j,k}$ | Probability or rate of a PRDM9 binding site with affinity $i$, recognized by PRDM9 allele $j$, experiencing gene conversion spanning the PRDM9 binding motif, in an individual with PRDM9 alleles $j$ and $k$. | |
| $g_i^j$ | Probability or rate of a PRDM9 binding site with affinity $i$, recognized by PRDM9 allele $j$, experiencing gene conversion spanning the PRDM9 binding motif, across the population. | |

## Results

### Fitness when considering one heat

Our model differs from previous ones (Úbeda and Wilkins 2011; Latrille et al. 2017; Úbeda et al. 2019) in assuming that: (i) individual fitness is a function of the probability that a DSB is made on the smallest chromosome at a site symmetrically bound by PRDM9, (ii) PRDM9 binding sites compete for a finite number of PRDM9 molecules, and (iii) PRDM9-bound sites compete for a constant number of DSBs. These last two assumptions are key, as can be understood by considering how fitness would change over time without them, under the simplest possible scenario, in which all individuals are homozygous for a single PRDM9 allele and all binding sites have the same binding affinity.

If we assume that there are always more PRDM9-bound sites than DSBs per meiosis, and ignore assumptions (ii) and (iii), such that the number of bound PRDM9 molecules is proportional to the number of binding sites and the number of DSBs is proportional to the number of bound PRDM9 molecules, fitness will decrease over time as binding sites are lost, as expected under Red Queen theories of hotspot evolution (**Fig 2A**: green line). This decrease in fitness is due to the dwindling numbers of DSBs per meiosis, however–the probability of symmetric binding per site remains constant (**Fig 2B**: blue lines). If instead, we assume that PRDM9-bound sites compete for a constant number of DSBs, as seems more realistic (Diagouraga et al. 2018), but continue to assume that PRDM9 binding is not competitive–i.e., ignore only assumption (ii)–fitness does not change over time, because both the number of DSBs per meiosis and the probability of symmetric binding per site remain constant (**Fig 2A**: blue line). Thus, there is no source of selection to drive the evolution of PRDM9. Lastly, if we consider a more biologically realistic model, in which there is competition for both PRDM9 binding and DSBs, fitness increases over time, with the loss of binding sites (**Fig 2A**: red line); indeed, as the number of binding sites decreases, the proportion of sites that are bound symmetrically increases (**Fig 2B**: red lines). Taken at face value, this model would suggest that selection should favor and hence retain older PRDM9 alleles, a prediction that is inconsistent with ubiquitous evidence for the rapid evolution of the PRDM9 ZF array’s binding specificity (Baker et al. 2017). As we discuss, this unrealistic behavior comes from considering only a single binding heat.

**Figure 2:**
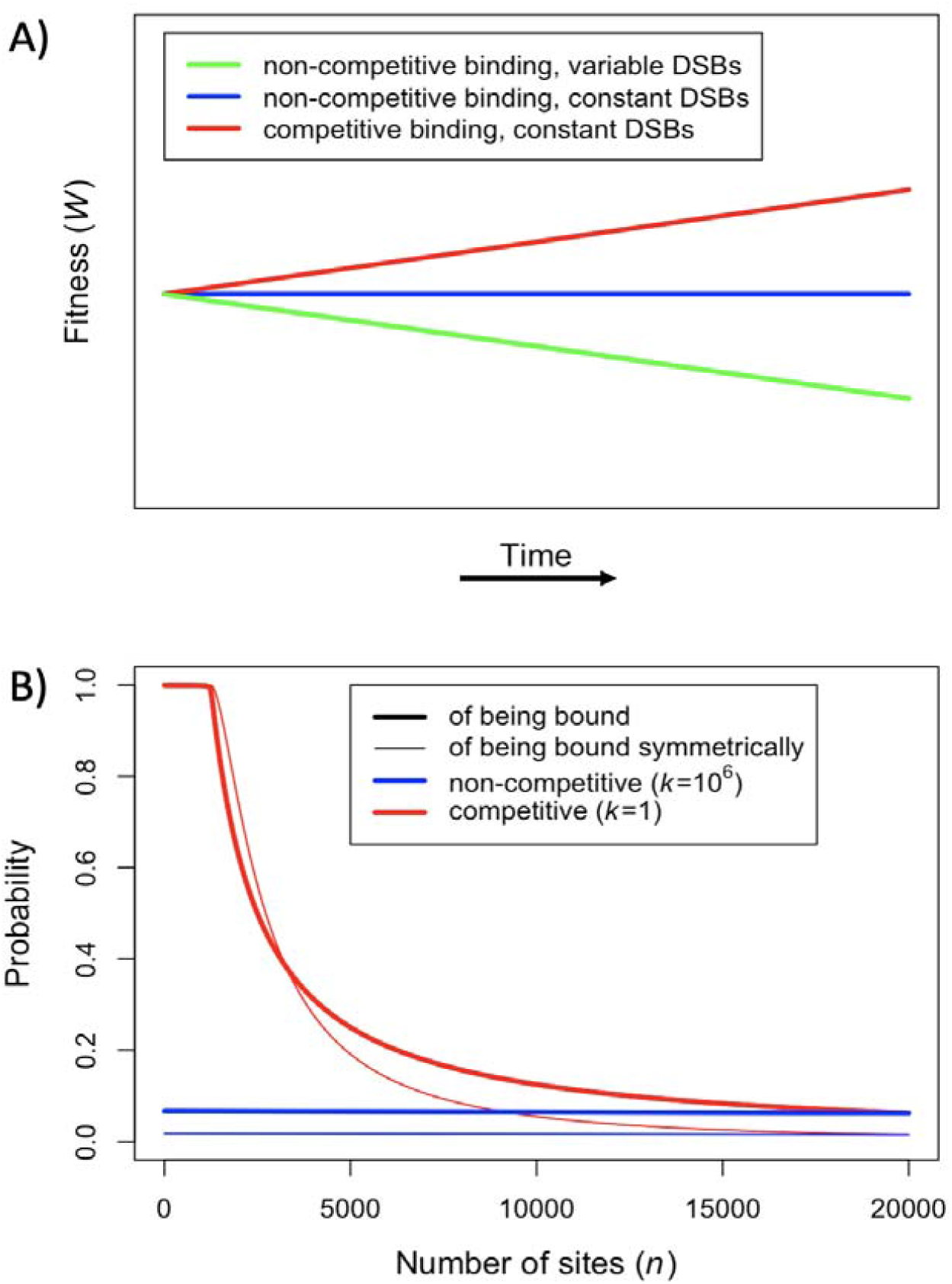
Fitness in the model with one heat. (**A**) Cartoon of fitness as a function of time under different assumptions. (**B**) The probability of an individual binding site being bound (thicker lines) or of a given locus being bound symmetrically (thinner line) if all sites are very weak (blue) or very strong (red), i.e., with very large or small dissociation constants respectively. For sake of comparison, in both cases, we set the number of expressed PRDM9 molecules such that 5,000 sites would be bound in the presence of 20,000 binding sites, roughly consistent with observations in mice (Parvanov et al. 2017; Grey et al. 2017). When binding sites are very weak, most PRDM9 molecules are not bound and therefore our model behaves as if there were no competition between binding sites.

### Fitness when considering two heats

To understand how fitness might decay in a more realistic setting, with competition among binding sites for PRDM9 molecules and among bound sites for DSBs, we turn to a model with two classes of binding sites. Specifically, we assume that there are a small number of strong binding sites that are bound symmetrically at substantial rates, and a larger number of weaker binding sites, which are not. Consequently, recombination occurs almost entirely at the strong binding sites, whose numbers decrease over time and determine how fitness behaves; the weak binding sites, whose numbers do not change substantially over time, provide the backdrop of competitive PRDM9 binding. For the sake of clarity, henceforth we refer to the stronger class of binding sites as ‘hotspots’ and the weaker class as ‘weak binding sites’.

The competitive effect of weak binding sites on individual hotspots is captured by the ratio of the dissociation constant at hotspots *k*_1_ to the proportion of PRDM9 bound in the absence of any hotspots (see **Supplementary Section S3**). We fix the number and dissociation constants of weak binding sites such that 99% of expressed PRDM9 molecules would bind the weak binding sites in the absence of hotspots, while contributing minimally to symmetric binding (*n*_2_ = 200,000, *k*_2_ = 8030). We then explore how the dynamics depend on the value of *k*_1_.

First, we consider how fitness depends on the number of hotspots, in individuals that are either homozygous or heterozygous for two PRDM9 alleles with the same binding distribution (**Fig 3**). Increasing the number of hotspots has conflicting effects on fitness: the proportion of bound PRDM9 localized to hotspots rises, and therefore there is a larger number of DSBs at hotspots, but the probability that any given hotspot is bound symmetrically decreases (**Fig 3 - Figure Supplement 1**). Increasing PRDM9 expression--and hence the number of PRDM9 molecules--has the opposite effects: it increases the probability that any given hotspot is bound symmetrically, but also increases the proportion of bound PRDM9, and thus DSBs, localized to weak binding sites (**Fig 3 - Figure Supplement 1**). Consequently, for any value of *k*_1_, there is an optimal number of hotspots, which maximizes the probability of forming symmetric DSBs, and this number increases with PRDM9 expression levels (in order to counteract PRDM9 localization to weak binding sites). These considerations imply that the optimal number of hotspots per allele is always smaller in heterozygotes relative to homozygotes (**Fig 3**). The optimal number of hotspots is also always smaller when hotspots are stronger, as fewer are needed to prevent PRDM9 binding to weak binding sites (**Fig 3**).

**Figure 3:**
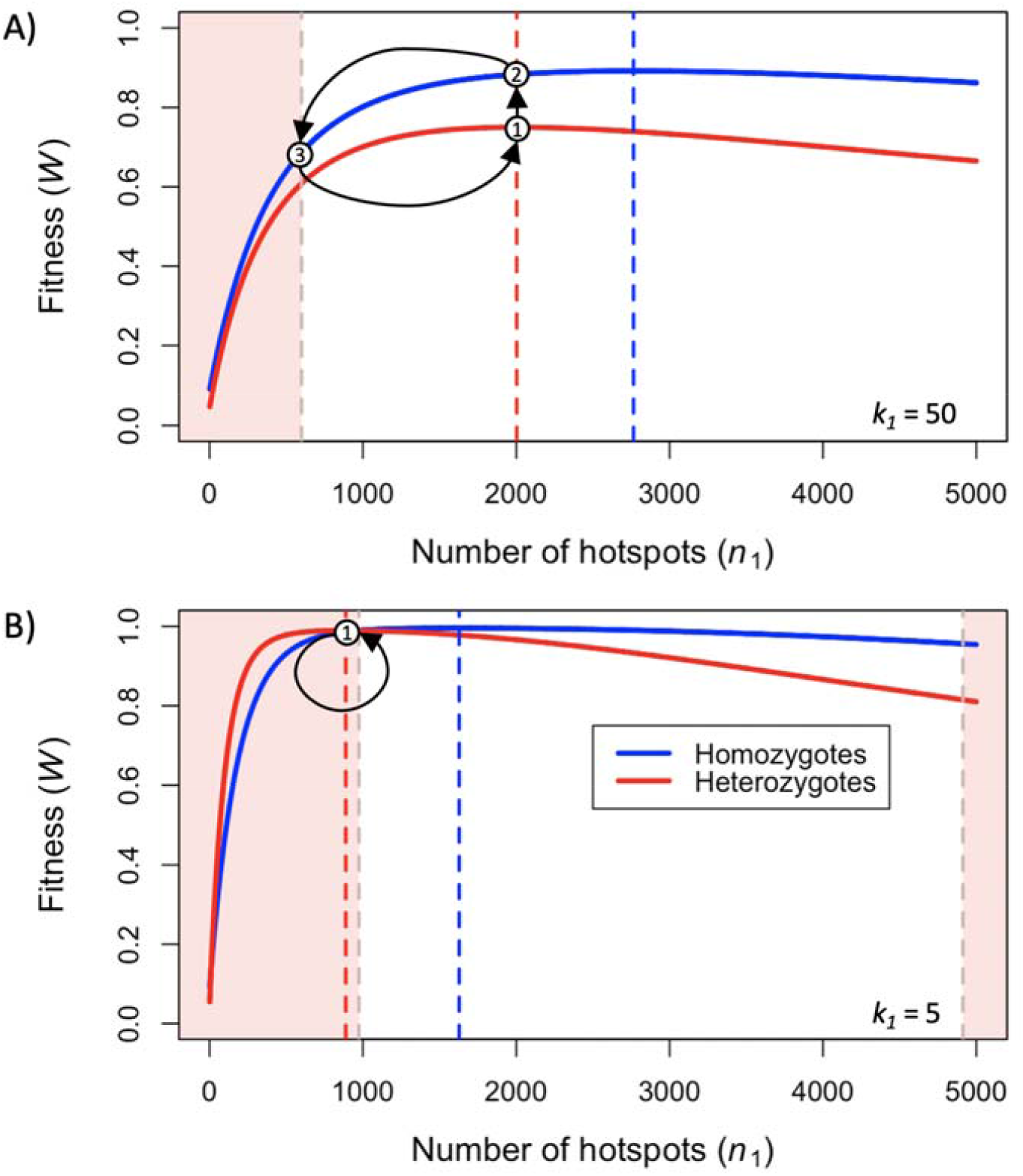
Fitness in the model with two heats. Fitness in an individual homozygous for a PRDM9 allele, or heterozygous for two PRDM9 alleles with the same binding distribution, when considering (**A**) weak hotspots or (**B**) strong ones. Vertical dashed lines indicate the number of hotspots that maximize the fitness of individuals homozygous (blue) or heterozygous (red) for the allele. The area shaded in light red indicates the number of hotspots of a PRDM9 allele where the fitness of an individual homozygous for the allele will have a lower fitness than the maximal fitness of an individual heterozygous for that allele (i.e., for that allele and one with the optimal number of hotspots for heterozygotes). The arrows between states (1)-(3) illustrate the behavior of the simplified evolutionary dynamic described in the text.

When hotspots are relatively weak (e.g., *k*_1_ = 50), fitness will always be higher in individuals homozygous for a given PRDM9 allele than in individuals heterozygous for two distinct PRDM9 alleles with the same number of hotspots (**Fig 3A**). Weak hotspots are far from being saturated for PRDM9 binding: the increased expression of a given PRDM9 allele in homozygotes relative to heterozygotes results in a notable increase in the probability that hotspots will be bound by PRDM9, while only slightly increasing the proportion of PRDM9 bound to weak binding sites (**Fig 3 - Figure Supplement 1A**). Consequently, the benefit of the increase in symmetric binding at hotspots outweighs the cost of the slight increase in the proportion of DSBs localized to weak binding sites.

In contrast, when hotspots are relatively strong (e.g., *k*_1_ = 5), fitness can be higher in individuals homozygous for a given PRDM9 allele or heterozygous for two similar alleles, depending on the number of hotspots recognized by the alleles (**Fig 3B**). When the number of hotspots far exceeds the number of PRDM9 molecules, hotspots are not saturated for PRDM9 binding, and fitness is always higher in homozygotes, as in the case with weaker hotspots. However, as hotspots are lost, those hotspots remaining approach saturation for PRDM9 binding and consequently, the increased expression of a given PRDM9 allele in homozygotes relative to heterozygotes has a lesser impact on rates of symmetric binding (**Fig 3 - Figure Supplement 1B**). Moreover, the cost associated with greater PRDM9 expression leading to an increased proportion of PRDM9 localizing to weak binding sites becomes more pronounced (**Fig 3 - Figure Supplement 1B**). The net effect is that, for PRDM9 alleles that recognize smaller numbers of hotspots, fitness is higher in heterozygotes than in homozygotes: the loss in fitness caused by PRDM9 localization to weak binding sites in homozygotes relative to heterozygotes outweighs the small increase in symmetric binding at hotspots. Thus, in contrast to the case for weaker hotspots, for stronger hotspots (e.g., *k*_1_ = 5), individuals heterozygous for two PRDM9 alleles, each with the optimal number of hotspots for heterozygotes, will have higher fitness than individuals homozygous for a PRDM9 allele with the same number of hotspots (**Fig 3B**).

The relationship between the fitness of homozygotes and heterozygotes provides important insights into the evolutionary dynamics of our model. Consider a simplified model, with a population of infinite size, in which mutations to PRDM9 alleles with any number of hotspots arise every generation. If individuals heterozygous for a given mutant allele have greater fitness than any other individual in the population, that allele will quickly invade the population and rise to appreciable frequencies. Under these conditions, the PRDM9 alleles that successfully invade the population will necessarily be those with optimal numbers of hotspots in heterozygotes. When hotspots are relatively weak (e.g., *k*_1_ = 50), the population exhibits cycles (**Fig 3A**): a PRDM9 allele with the optimal number of hotspots in heterozygotes invades the population and eventually fixes (arrow 1 in **Fig 3A**), after which individuals are homozygous for that allele and have greater fitness than any PRDM9 heterozygote, thereby preventing any other new allele from invading the population. Hotspots will then be lost from the population (arrow 2 in **Fig 3A**), until another PRDM9 allele can invade (arrow 3 in **Fig 3A**), fix and begin a new cycle. When hotspots are relatively strong (e.g., *k*_1_ = 5), the population remains at a fixed point at which PRDM9 alleles with the optimal number of hotspots in heterozygotes are constantly invading (**Fig 3B**): once such an allele reaches appreciable frequencies and individuals become homozygous for the allele, their fitness becomes lower than that of a heterozygote for a new PRDM9 alleles with the optimal number of hotspots, and this prevents from them reaching fixation. These two kinds of behaviors, of cycles and a fixed point, carry over with some modifications to the case of more complex models.

### Dynamics of the two-heat model

Next, we consider the dynamic in a finite population size and given a mutation rate at the PRDM9 locus. We assume that the number of hotspots associated with new PRDM9 alleles is uniformly distributed across a wide range (*S*_1_ = 1 − 5000), which spans the optimal values for both homozygotes and heterozygotes across the range of hotspot dissociation constants that we explore (*k*_1_ = 5 − 50). Where possible, the values that we use for model parameters are based on empirical observations, including the mutation rates at the PRDM9 locus and its binding sites, the number of DSBs initiated per meiosis, and the probability that a gene conversion event removes the hot allele at a PRDM9 binding site in individuals heterozygous for hot and cold alleles (**Table 1**). We focus on how the evolutionary dynamics depend on the dissociation constant at hotspots, *k*_1_, which we know less about, and the effective population size, which varies over orders of magnitude in natural populations (Leffler et al. 2012).

When the population size is large (e.g., *N* = 10^6^), the dynamics are similar to that of the infinite population size case (**Fig. 4A** and **C**). When hotspots are fairly weak (e.g., *k*_1_ = 50), they follow an approximate cycle (**Fig 4A**): a new PRDM9 allele that establishes a near optimal number of hotspots for heterozygotes invades and fixes, dominating the population in a homozygous state until enough hotspots are lost that the population becomes susceptible to invasion of a new allele, likewise with a near optimal number of hotspots for heterozygotes. Diversity at the PRDM9 locus is low throughout most of this cycle, with bouts of diversity in the period during which the dominant allele is taken over by a new one. Fitness peaks when a new allele is fixed and then decreases as that allele loses its strong binding sites.

**Figure 4.**
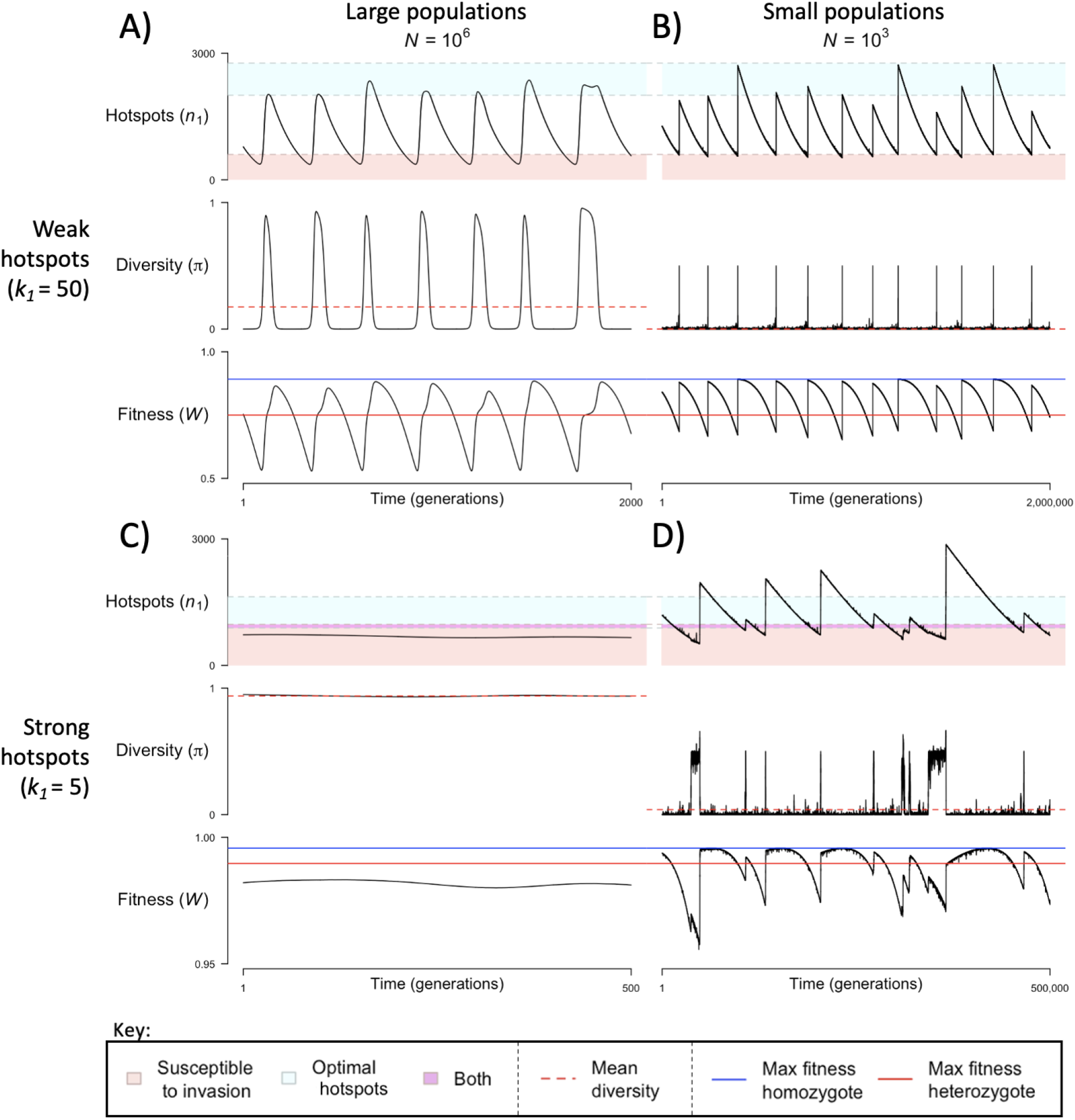
The dynamics of the model with two heats. The mean number of hotspots () amongst segregating alleles, diversity at PRDM9 (), and mean fitness (), as a function of time. We show four cases: with weak and strong hotspots (top and bottom rows respectively) and large and small population size (left and right column respectively). The time (x-axis) in each plot has been scaled by the duration of PRDM9 turnover to allow for ~10 ‘cycles’ (see **Fig. 3**), which span ~10^3^ times more generations with the smaller compared to the larger population size. The range of hotspots ‘susceptible to invasion’ shown correspond to the regions shaded in red in **Fig 3**. The range of ‘optimal hotspots’ corresponds to the number that optimizes fitness in heterozygotes to the number that optimizes fitness in homozygotes.

In contrast, when hotspots are strong (e.g., *k*_1_ = 5), dynamics are approximately at a fixed point (**Fig 4C**): new PRDM9 alleles that lead to near optimal numbers of hotspots for heterozygotes continually invade but never fix. The fitness of individuals homozygous for such PRDM9 alleles is lower than that of individuals heterozygous for one such allele and those already present. Consequently, new PRDM9 alleles experience a kind of frequency-dependent selection, in which they are favored at low frequencies because they have near optimal numbers of hotspots for heterozygotes, but selected against at higher frequencies because of the reduced fitness of homozygotes. As a result, diversity remains high and average fitness is approximately constant.

In smaller populations (e.g., *N* = 10^3^), the dynamic is qualitatively similar to that in larger populations when hotspots are fairly weak (e.g., *k*_1_ = 50; compare **Fig 4A** and **B**). The reduced mutational input at hotspots and correspondingly lower rate at which they are lost leads to longer intervals of time between fixation events. In turn, the reduced mutation input at PRDM9 and stronger drift leads to reduced peaks of PRDM9 diversity during those events.

In contrast, when hotspots are strong (e.g., *k*_1_ = 5), dynamics differ markedly in smaller population compared to larger ones (compare **Fig 4C** and **D**): whereas in large populations, PRDM9 alleles never fix, in small populations, the lower mutational input at the PRDM9 locus and stronger genetic drift allows PRDM9 alleles to fix intermittently (**Fig 4D**). PRDM9 alleles with a small number of hotspots are typically lost rapidly and without reaching fixation--the same behavior as in larger populations. But when a mutation occurs to a PRDM9 allele with a large number of hotspots, such that homozygotes for it have higher fitness than any heterozygotes (see **Fig. 3B**), and the allele happens to reach high frequency by chance, it can be maintained in the population at appreciable frequencies–until it has lost a sufficient number of hotspots for the population to become susceptible to the invasion of a new PRDM9 allele. These chance fixation events and the corresponding number of hotspots leads to intermittent, chaotic cycles.

In both small or large populations, stochasticity in the PRDM9 mutations that arise and escape immediate loss gives rise to a distribution of initial number of hotspots associated with segregating PRDM9 alleles. We obtain this distribution by weighting PRDM9 alleles by the product of their sojourn times and mean frequencies (**Fig 5**). This distribution is centered close to but above the optimal number of hotspots for heterozygotes, in part because newly arising alleles are always found in heterozygotes (**Fig 5** and **6A**). The mean is generally closer to this optimal number in large populations than in small populations, reflecting the greater mutational input at PRDM9 (**Fig 6A**). However, when hotspots are strong and the population size is small, the intermittent fixations of PRDM9 alleles with a large number of hotspots shift the distribution towards larger initial numbers of hotspots (**Fig 5** and **Fig 6A**).

**Figure 5.**
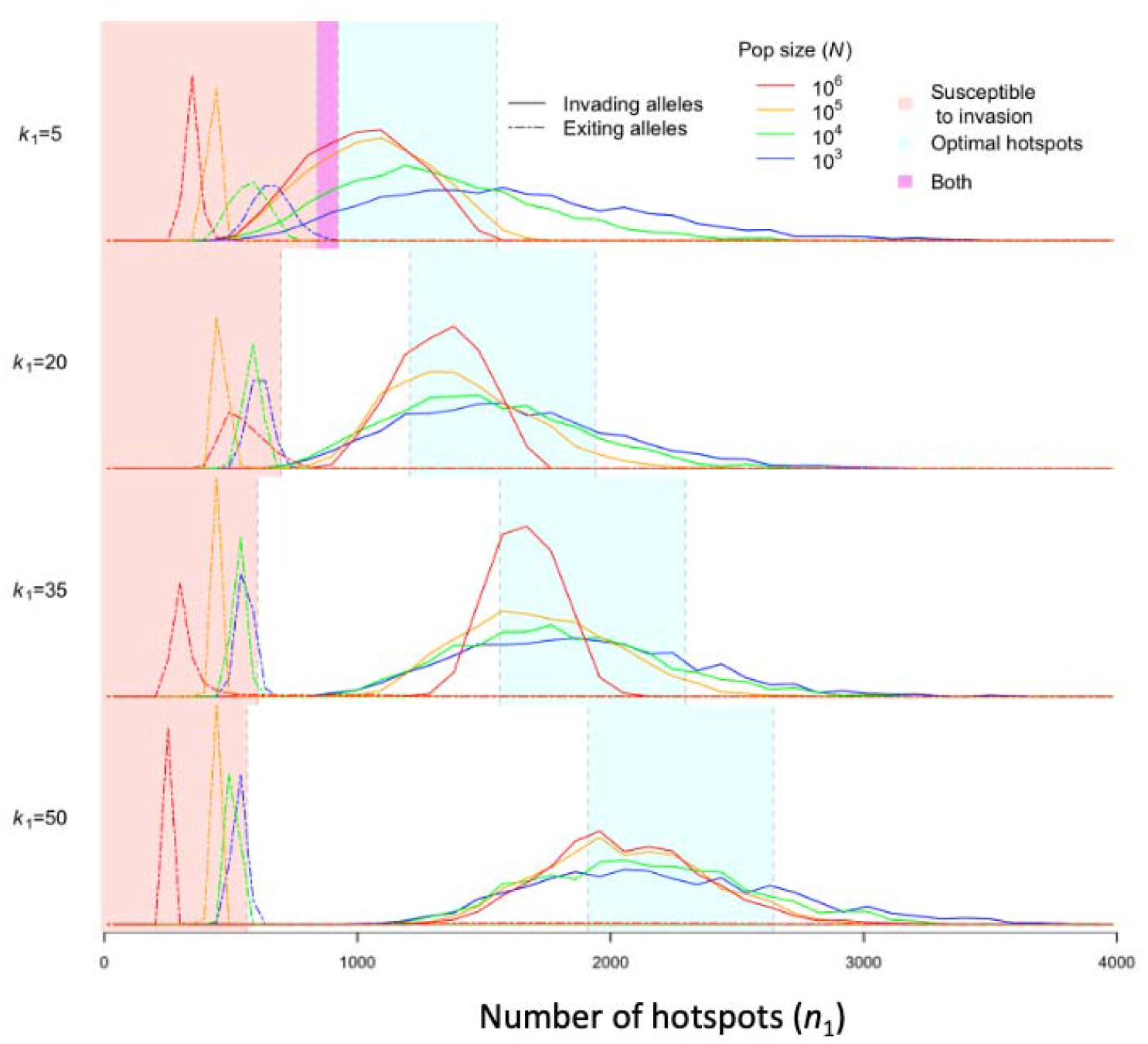
The distribution initial and final numbers of hotspots among segregating PRDM9 alleles. The relative densities of initial and final numbers of hotspots for different dissociation constants (rows) and population sizes (line colors) obtained from simulations of five million PRDM9 alleles (see text).

**Figure 6.**
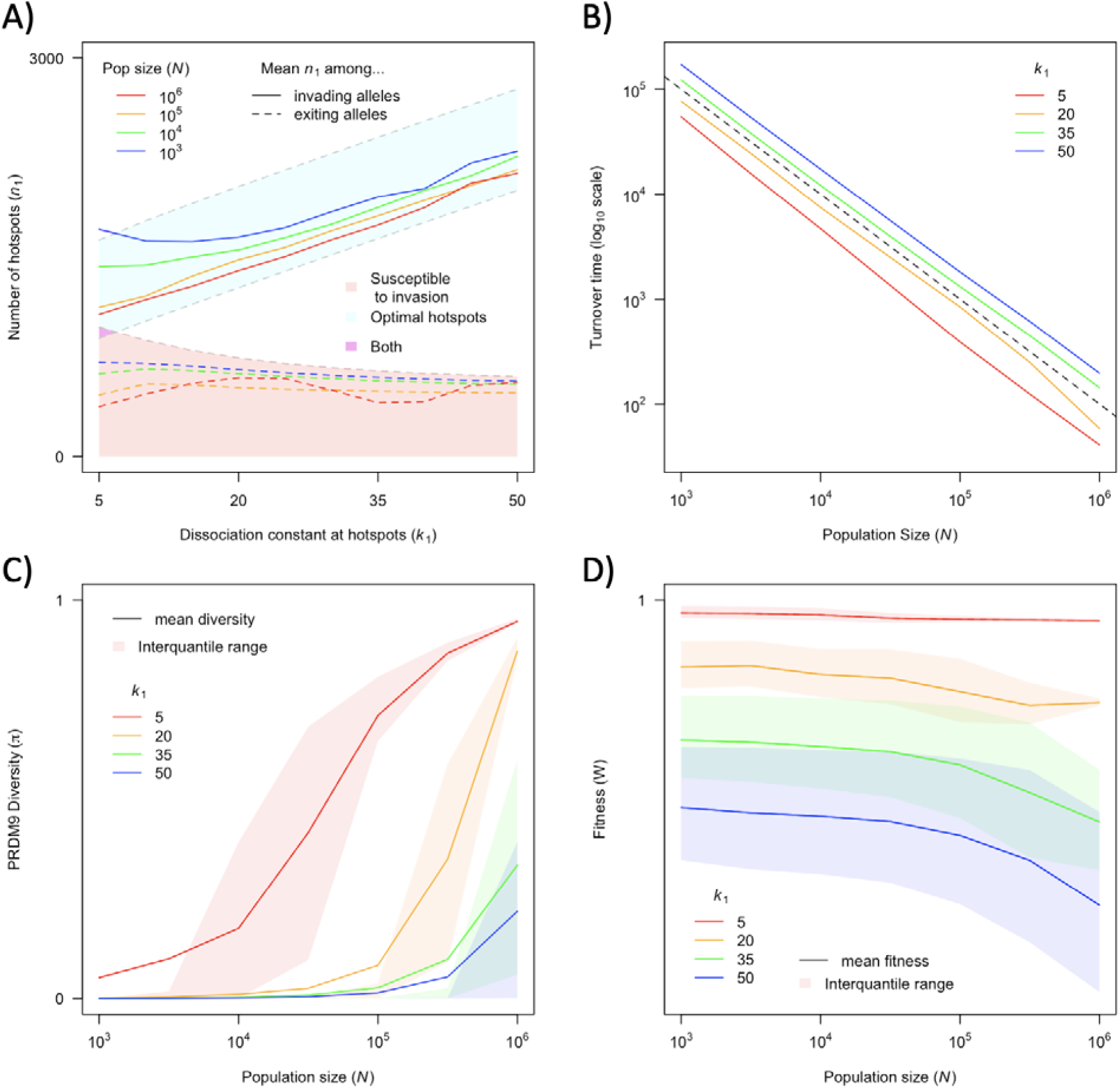
The dependence of key quantities on population size and hotspot strength. (**A**) Mean number of hotspots among incoming and exiting alleles (corresponding to the distributions shown in **Fig 5**). (**B**) Average turnover time of segregating PRDM9 alleles (see text). (**C**) Mean diversity at PRDM9 over time. (**D**) Mean fitness in the population.

In turn, the distribution of the final numbers of hotspots associated with PRDM9 alleles exiting the population (similarly weighted by sojourn times and mean frequencies) is centered below the number at which they become selected against (at which the population fixed for such an allele would become susceptible to invasion for one with the optimal numbers of hotspots for heterozygotes), because they continue to lose hotspots during the time it takes a new PRDM9 allele to fix. More hotspots are lost during this time interval in larger populations than in smaller populations, owing to the increased mutational input at hotspots (**Fig 5** and **6A**).

The turnover time at PRDM9, defined as the mean product of sojourn times and mean frequencies across PRDM9 alleles, is determined principally by the rate at which hotspots are lost and how many hotspots need to be lost on average before an allele becomes selected against. In particular, turnover time is inversely proportional to population size (**Fig 6B**). This relationship is driven almost entirely by the increased mutational input at PRDM9 binding sites in larger populations (and correspondingly increased rate at which hotspots are lost), while the mutational input at PRDM9 is only minorly rate-limiting in smaller populations (**Fig 6 - Figure Supplement 1A, Fig 6 - Figure Supplement 2**). Similarly, the turnover time is reduced for strong hotspots relative to weak one (**Fig 6B**). This reduction results from the lower initial number of hotspots (**Fig 6A**), as well as from the fact that stronger hotspots experience more DSBs and are correspondingly lost more quickly.

The effects of the population size and the dissociation constant of hotspots on turnover times shape diversity levels at PRDM9. Inversely, the peak levels of diversity observed are influenced primarily by the mutation rate at PRDM9 (**Fig 6 - Figure Supplement 2**, compare rows 1 and 2). The effect of population size on PRDM9 diversity is explained in part by the mutational input at binding sites decreasing the time between turnover events (**Fig 6 - Figure Supplement 2**, compare rows 1 and 3), and the high levels of diversity that accompany them, and in part by the mutational input at PRDM9 increasing diversity during such periods (**Fig 6C, Fig 6 - Figure Supplement 1B**). As explored above, the strength of hotspots additionally plays an important role in shaping patterns of PRDM9 diversity by determining the relative fitnesses in heterozygous or homozygous individuals for different PRDM9 alleles, and thus whether alleles will typically have a homozygous or heterozygous advantage.

Notably, the co-evolutionary dynamics of PRDM9 and its binding sites entail a substantial genetic load, i.e., the average fitness is substantially lower than the optimal value. Moreover, and in contrast to the usual tenet of population genetics, the average fitness is greater in smaller populations (**Fig 6D**). These phenomena arise for two reasons. First, the greater efficacy of selection and mutational input at PRDM9 in larger populations shifts the distribution of initial numbers of hotspots among invading alleles towards values optimizing fitness in heterozygotes (**Fig 5** and **6A)**. Inversely, because the maximum possible fitness for homozygous individuals is always higher than that of heterozygotes (**Fig 3**), the optimal number of initial hotspots for maximizing fitness across the population will be closer to that which optimizes fitness in homozygotes. Because the shift towards optimal values for heterozygotes tends to bring those values further from what would be optimal for homozygotes, this tends to cause a reduction in mean fitness. Second, the increased mutational input at hotspots in larger populations leads to lower (and less optimal) numbers of hotspots recognized by PRDM9 alleles segregating in the population at any given time (**Fig 5** and **6A**).

## Discussion

Previous models of PRDM9 evolution demonstrated how selection to maintain a given rate of recombination (e.g., for there to be at least one crossover per chromosome) can act as a countervailing force to the loss of hotspots over time through gene conversion and ensure the persistence of hotspots (Úbeda and Wilkins 2011; Latrille et al. 2017). These models were based on an assumption now believed to be incorrect, however: that the loss of PRDM9 binding sites leads to a reduction in DSB formation. Notably, DSB formation is regulated independently of PRDM9 (Kauppi et al. 2013; Yamada et al. 2017; Brick et al. 2012; Mihola et al. 2019) and hotspots are thought to compete for binding (Diagouraga et al. 2018); when hotspots are lost, PRDM9 may simply bind elsewhere. Perhaps most relevant, recent evidence indicates that the successful completion of meiosis requires the formation of DSBs at sites symmetrically bound by PRDM9 across homologous chromosomes (Smagulova et al. 2016; Davies et al. 2016; Gregorova et al. 2018; Hinch et al. 2019; Li et al. 2019; Huang et al. 2020; Mahgroub et al. 2020; Wells et al. 2020; Gergelits et al. 2021; Davies et al. 2021).

Motivated by these discoveries, we developed a new model for the evolution of PRDM9, considering the competition of binding sites for PRDM9 molecules and of PRDM9-bound sites for DSBs, in which the loss of individual binding sites leads to increased PRDM9 binding and DSB formation at remaining sites. In this setting, the rates at which binding sites are lost from the population due to GC and the probabilities that they experience symmetric DSBs are each determined by their underlying affinities for PRDM9. Strong hotspots are lost more rapidly than weak ones, resulting in a shift from a small number of strong binding sites to a larger number of weak binding sites over the lifespan of a PRDM9 allele. This shift is accompanied by a reduction in the number of symmetric DSBs, and increasing selection against that PRDM9 allele. New PRDM9 alleles restore binding sites, thereby serving as a countervailing force to GC. But they are favored because, in doing so, PRDM9 binding is redirected to a smaller proportion of the genome, and correspondingly is more likely to bind symmetrically. Accordingly, we suggest that the selection driving the evolution of the PRDM9 ZF arises from the requirement of symmetric binding in DSB repair, perhaps ultimately because of homolog pairing, and thus from changes in the shape of its binding distribution over time, rather than from changes in net binding, as previously believed.

We employed a number of simplifying assumptions in our model. Notably, we assumed a very simple distribution of heats that suffices to demonstrate key behaviors of PRDM9 evolution, but is likely insufficient to capture all quantitative dynamics. Considering a more complete distribution (i.e., with many heats) may be necessary to more accurately characterize the timescale of PRDM9 turnover, for example, or how the fitness of an allele in heterozygotes changes over time relative to homozygotes. We also modeled the loss of hotspots deterministically, without considering the contribution to asymmetry that might stem from heterozygosity at binding sites for instance. Also not considered in the model is the possibility that when PRDM9 is not bound symmetrically, occasional repair off the sister chromatid rather than the homolog could reduce the rate at which hotspots are lost, while a mutagenic effect of recombination may increase the rate at which hotspots are lost. Lastly, by assuming that the majority of PRDM9 is bound in the absence of strong binding sites, and that all PRDM9-bound sites are equally likely to experience a DSB, we implicitly assumed that each allele in heterozygotes was responsible for roughly half of all PRDM9-bound sites, and half the DSBs per meiosis, thereby ignoring some potential dominance effects. Despite these limitations, our model already provides important new insights into PRDM9 evolution and the selection pressures that shape the recombination landscape, as discussed below.

### The requirement for symmetry shapes the recombination landscape

In our two-heat model, PRDM9 alleles can arise with different binding distributions (i.e., with different numbers of hotspots), and selection favors PRDM9 alleles for which PRDM9 binding is concentrated to the smallest proportion of the genome (i.e., with a relatively small number of hotspots, but enough to prevent PRDM9 binding to the more numerous weaker sites). In a more realistic model, with many heats, we would likewise expect selection to favor highly skewed distributions, i.e., with hotspots, as is observed empirically.

The benefit of PRDM9 binding symmetry presumably lies in the fact that the same locus used for DSB initiation on one chromosome is, on the homolog, preferentially used as template for DSB repair. As far as we are aware, Davies et al. 2016 was the first to hypothesize that this might reflect a more general benefit of using hotspots for localizing meiotic recombination: that preferentially using a small proportion of the genome as both sites for DSB initiation and templates for DSB repair vastly reduces the search space during homolog engagement, i.e., the number of genomic positions each DSB needs to be checked against before finding the appropriate template for repair (Davies et al. 2016). A number of subsequent studies have provided further support for this hypothesis as it pertains to PRDM9 (Gregorova et al. 2018, Hinch et al. 2019; Li et al. 2019; Gergelits et al. 2020; Huang et al. 2020, Mahgroub et al. 2020, Wells et al. 2020). In particular, hotspots that are more likely to be bound symmetrically are repaired more rapidly (Hinch et al. 2019; Li et al. 2019).

In our model, we considered individual fitness to be the probability that at least one DSB forms on the smallest chromosome at a site that had been symmetrically bound by PRDM9. In doing so, we considered only the probability that at least one site would find the appropriate template for repair after preferentially checking each of *n* other PRDM9-bound sites. A more complex model might further consider that the benefit of symmetry is a function of *n*, i.e., that the benefit of symmetry would be improved if even fewer sites needed to be checked, or inversely, that being symmetrically bound would be less predictive of homology in the presence of a very large number of bound sites. Although we do not explicitly consider changes in the size of the search space that result from changes in net PRDM9 binding, our results suggest an implicit benefit to the use of a smaller number of sites for recombination, when considering competition between binding sites for PRDM9 and between bound-sites for DSBs (as shown in **Fig. 1A**).

It further stands to reason that if binding symmetry, in terms of PRDM9 or of downstream factors involved in DSB initiation and repair, enables DSB repair, it also likely serves to improve the efficacy of DSB-dependent mechanisms of homolog pairing and synapsis. In this regard, it is interesting that many taxa with DSB-independent mechanisms of homolog pairing, such as *Drosophila* or *C. elegans*, do not have hotspots, whereas most species with DSB-dependent mechanisms, such as yeast, plants or vertebrates, do. We speculate that the general prevalence of hotspots may be related to the distribution of DSB-dependent mechanisms for proper homolog pairing and segregation.

### Does the decay of hotspots by GC lead to more or fewer hotspots?

Intriguingly, the loss of individual hotspots in our model may lead to an increase or a decrease in the total number of hotspots in our model, depending on the distribution of heats considered, as well as how hotspots are defined. The standard definition of hotspots, also used operationally to identify them, is as short intervals of the genome (typically < 2 kilobases) with recombination rates that exceed the local or genome-wide background rate by more than some arbitrary multiplicative factor (e.g., five-fold; reviewed in Tock and Henderson 2018). In the two-heat model that we explored, the stronger class of binding sites would easily be identified as hotspots, whereas the weaker binding sites would be taken as non-specific binding, if they were detected at all. However, if we were to consider more realistic binding distributions, we would find some binding sites with heats just below any given threshold used to define hotspots, which would become hotspots after some of the stronger ones are lost. Given such a distribution, we could define a threshold such that the number of hotspots increases over the lifespan of a PRDM9 allele as its binding becomes more dispersed along the genome.

We liken this observation to the optical illusion of the Penrose stairs. Walking around the stairs in one direction, there is a gradual decrease in the number of symmetrically-bound sites over time, that is suddenly increased upon the appearance of a new PRDM9 allele. Walking in the other direction, there is a gradual increase in the number of loci that are used for recombination, that is suddenly decreased upon the appearance of a new PRDM9 allele. Either way you go around, however, you end up in the same spot.

### Drive against most binding sites is not an important driver of PRDM9 evolution

The loss of individual PRDM9 binding sites over time occurs because GC effectively acts as selection against hot alleles. The strength of this selection in our two-heat model (*s* ≈ *0.006* to 0.021, corresponding to the range of dissociation constants considered) is greater than that used in previous models of PRDM9 evolution (*s* ≈ 0.003; Latrille et al. 2017), but far weaker than can be inferred empirically from the under-transmission of exceptionally hot alleles within humans (*s* ≈ 0.46 to 0.53; Jeffreys and Neumann et al. 2002; Berg et al. 2011). Nonetheless, more recent estimates of global rates of drive at PRDM9 binding sites have suggested that the strength experienced by most hotspots is even weaker than previous modeling work assumed (*s* ≈ 5 × 10^−5^ to 2 × 10^−4^; Spence and Song 2019). These empirical estimates (while plausibly downwardly biased; Spence and Song 2019) suggest that the strength of drive against most PRDM9 binding sites may be on the same order as the inverse of the effective population size, and thus that this drive is a weak evolutionary force. In light of these results, Spence and Song (2019) suggested that the evolution of PRDM9 might be explained if only a small subset of PRDM9 binding sites were crucial for the proper segregation of chromosomes during meiosis and if the drive against these hotspots was much stronger. This conjecture is borne out by our model, in which both the rate of DSB formation and the probability of being bound symmetrically by PRDM9 are determined by the underlying binding affinity of a given site. Accordingly, the sites most likely to be symmetrically bound will also experience the highest rates of drive, in accordance with empirical observations (Hinch et al. 2019). We therefore suggest that the hotspots in the two-heat model can be thought of as approximating the behavior of the strongest PRDM9 binding sites within a more realistic, continuous distribution of binding affinities.

### Diversity of the PRDM9 ZF-array

As in previous models of PRDM9 evolution (Latrille et al. 2017), we expect diversity levels reported by our model should generally be lower than is observed empirically, both because population structure contributes to observed diversity (and we assumed a panmixia), and because naturally-occurring PRDM9 alleles will sometimes share some overlap of binding sites (and would be considered the same allele in our model).

PRDM9 diversity appears to be relatively high whether the effective population size is small or large (Buard et al. 2014; Vara et al. 2019; Alleva et al. 2021). Like all loci, PRDM9 diversity reflects the interplay between mutation, selection and genetic drift. While the rate of mutation at the PRDM9 ZF array has been inferred to be high (Jeffreys et al. 2013), as might be expected from its minisatellite nature, this high mutation rate may be counterbalanced by strong selection favoring specific PRDM9 ZF alleles (Úbeda and Wilkins 2011; Latrille et al. 2017; this study). In our model, the balance between mutation, selection and drift, leads to appreciable diversity only in large populations, where it is driven by the increased mutational input both at PRDM9 and at PRDM9 binding sites. Our model also suggests that diversity at PRDM9 could be enhanced by negative frequency-dependent selection that arises as a consequence of particular binding distributions. However, this result could reflect the large disparity between heats in our two-heat model, and it remains to be seen if the same behavior arises with more realistic distributions of binding affinities.

### Why use PRDM9?

PRDM9 arose prior to the common ancestor of animals and has since been repeatedly lost, including more than a dozen times across vertebrates (Baker et al. 2017; Cavissim et al. 2022). In vertebrate lineages in which PRDM9 has been lost, such as canids, birds or percomorph fish, as well as in PRDM9 knockout mice, recombination events are concentrated in hotspots that coincide with promoter-like features of the genome, such as transcriptional start sites and/or CpG-islands (Brick et al. 2012; Auton et al. 2013; Singal et al. 2015). The same is true for lineages that are not believed to have ever carried PRDM9 but still have recombination hotspots, such as in plants or fungi (Yelina et al. 2015; Lam and Keeney 2015). Therefore, this recombination mechanism is likely ancestral to the evolution of PRDM9’s role in recombination, and still conserved in the presence of PRDM9. How and why PRDM9 is repeatedly lost remains unknown however, as is the benefit PRDM9 typically confers over the use of default hotspots.

The selection that drives PRDM9 evolution in our model may help answer the latter. If the benefit of hotspots in species without PRDM9 is that they too allow for the recruitment of factors involved in both DSB initiation and DSB repair on both homologs, then PRDM9 may be favored when it further enhances the degree of symmetric recruitment of such factors relative to that of default hotspots, i.e., when PRDM9-mediated hotspots are fewer and hotter than default hotspots given the same number of DSBs. In support of this possibility, DSB heats measured by DMC1 ChIP-seq are higher in wild type than in PRDM9-/- mice (Brick et al. 2012).

If PRDM9 does indeed improve the efficiency of DSB-dependent homolog engagement, it may have the secondary benefit of enabling the successful completion of meiosis with fewer DSBs. While the number of DSBs formed per meiosis is regulated independently of PRDM9, we speculate that, given a requirement for a certain number of ‘symmetric DSBs’ per meiosis, the number of DSBs made in a typical germ cell should co-evolve with the degree of symmetric PRDM9 binding. Such a mechanism could explain the observation that not all strains of mice are sterile when PRDM9 is knocked out, and those that are tend to make fewer DSBs per meiosis (Mihola et al. 2019).

Thus, our model suggests a possible benefit of adopting PRDM9, in further improving the efficacy of the recombination process. Another evolutionary puzzle remains open, however: why PRDM9 has been lost so many times in different lineages (Baker et al. 2017, Cavassim et al. 2022). It is tempting to speculate that the answer to this question also relates to the advantage of symmetry. Perhaps in these species, segregating PRDM9 alleles no longer provided an advantage over the default use of promoter-like features, possibly because the use of such features in those species result in more symmetry than in others.

## Acknowledgements

We would like to thank Nicolas Lartillot, Laurent Duret, Scott Keeney, and members of the Przeworski and Sella labs for helpful discussions. This work was supported by NIH grant R01 GM83098 to MP and NIH R01 GM115889 to GS.

## Supplement

### Supplementary Section 1 – Molecular dynamics of PRDM9 binding

We first consider the case in which all binding sites for a given PRDM9 allele have the same binding affinity. At equilibrium, the rate at which PRDM9 molecules dissociate from their binding sites equals that rate at which they bind them. Denoting the expected number of bound PRDM9 molecules by *P*_*B*_ and the rate of dissociation of a bound PRDM9 molecule by *k*_*off*_, the total rate of PRDM9 dissociation is *k*_*off*_ · *P*_*B*_ (see Table1 for a summary of notation). Further denoting the expected number of free PRDM9 molecules by *P*_*F*_, the expected number of free binding sites by *n*_*F*_ and the rate of association between a free PRDM9 molecule and a given binding site by *k*_*on*_, the total rate at which PRDM9 molecules bind is *k*_*on*_ · *n*_*F*_ · *P*_*F*_. Therefore, at equilibrium

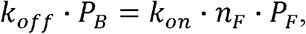

or equivalently

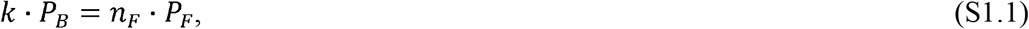

where *k* ≡ *k*_*off*_/*k*_*on*_ is the dissociation constant. Substituting *n*_*F*_ = 4*n* − *P*_*B*_ (where n is the total number of binding sites *per chromatid*) into Eq. S1.1 and solving for the expected number of bound PRDM9 molecules, we find that

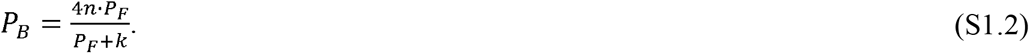

In this way, we see that the probability that a given site is bound by PRDM9, its “heat”, is

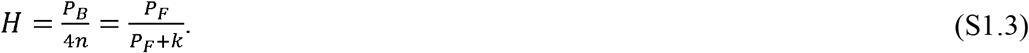

Next, we solve for the expected number of free PRDM9 molecules, *P*_*F*_. Substituting *P*_*B*_ = *P*_*T*_ − *P*_*F*_ (where *P*_*T*_ is the total number of PRDM9 molecules) into Eq. S1.2 and rearranging it, we end up with a quadratic in *P*_*F*_:

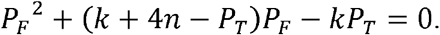

The general solution for this quadratic is

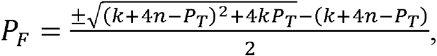

but further noting that 4*kP*_*T*_ >0 and that the expected number of free PRDM9 molecules must be positive, we find that

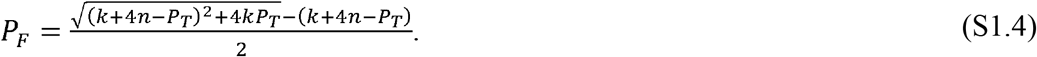

The case in which a PRDM9 molecule has binding sites with different affinities can be treated analogously. In this case, the equilibrium requirement must be met for sites of each affinity. By analogy to Eq. S1.2, we find that

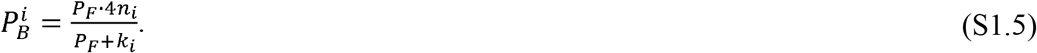

where 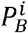 is the expected number of PRDM9 molecules bound to sites with the *i*-th affinity, *n*_*i*_ is the total number of these sites and *k*_*i*_ is their dissociation constant. Similarly, by analogy to Eq. S1.3, we find that the probability that a binding site with the *i*-th affinity is bound by PRDM9, its “heat”, is

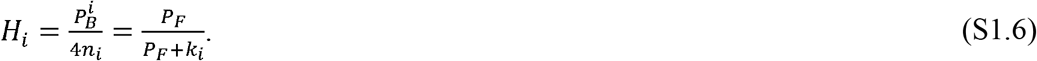

In order to solve for the expected number of free PRDM9 molecules, we note that

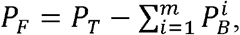

where *m* is the number of different binding affinities. Substituting the expression for 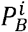 from Eq. S1.5 into this one, we find that

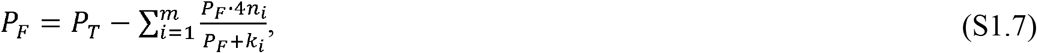

which is a polynomial of degree *m* + 1 in *P*_*F*_ that can be solved numerically.

In the case with two binding affinities, rearranging Eq. S1.7 yields the cubic equation:

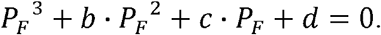

with coefficients *b* = (4*n*_1_ + 4*n*_2_ + *k*_1_ + *k*_2_ − *P*_*T*_), *c* = (4*k*_1_ *n*_2_ + 4*k*_2_ *n*_1_ + *k*_1_*k*_2_ − (*k*_1_ + *k*_2_) *P*_*T*_) and *d* = −*k*_2_*k*_1_*P*_*T*_. The solution of this equation is given by

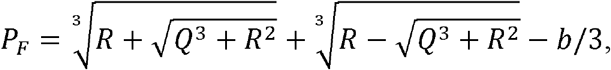

where *R* = (9*bc* − 27*d* − 2*b*^3^)/54 and *Q*= (3*c* − *b*^2^)/9.

### Supplementary Section 2 – The probability of a symmetric DSB

We define a “symmetric DSB” at a given binding site as the event in which, considering the four chromatids, at least one of four copies of the binding site (across the four copies of the chromosome present) during meiosis has been bound by PRDM9 and experienced a DSB and at least one of the two copies on the homologous chromosomes has also been bound by PRDM9.

We first consider the probability that at least one of the two copies on a pair of sister chromatids has been bound by PRDM9 and experienced a DSB. The probability that a given copy with binding affinity *i* has been bound is *H*_*i*_ ; having been bound, the probability that it experiences a DSB is *c*, and therefore the probability of both is *cH*_*i*_. The probability that at least one of the two copies on sister chromatids experienced both is the complement of the probability that neither of them did, that is

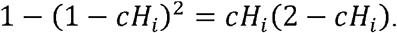

Similarly, the probability that at least one of the copies on the homologous chromosomes has been bound by PRDM9 is

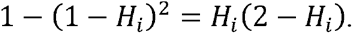

Assuming independence, the probability that the binding site on a given homologous chromosome has been bound and experienced a DSB and that the corresponding site on the other homologous chromosome has been bound is the product of the above expressions

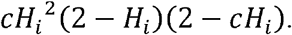

Lastly, given that either chromosome could be the one to experience the DSB, the probability of a ‘symmetric DSB’ is the complement of the probability that a ‘symmetric DSB’ occurs in neither direction, that is

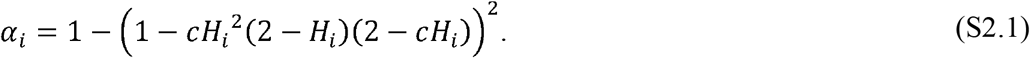

By the same token, the probability that a site with binding affinity *i* has been symmetrically bound, regardless of DSBs (as shown in **Fig. 2B**), is the complement of the probability that that neither chromatid on either homolog has been bound, i.e.,

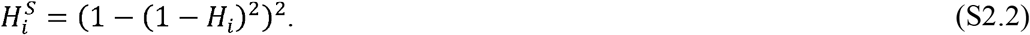

### Supplementary Section 3 – The competitive effect of weak binding sites in the two-heat model

The molecular and evolutionary dynamics of a PRDM9 allele in the two-heat model depends on three parameters that describe allele properties and on two variables. The parameters are the number of PRDM9 molecules, *P*_*T*_, the dissociation constant of hotspots, *k*_1_, and the dissociation constant of weak binding sites, *k*_2_; the variables are the total numbers of hotspots, *n*_1_, and of weak binding sites, *n*_2_. Here we show that in the parameter regime of interest to us, we can describe the dynamics in terms of a single variable 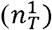 and two parameters: *P*_*T*_ and 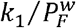, where 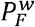 is the expected number of free PRM9 molecules in the absence of hotspots. The latter, compound parameter reflects the competitive effect of the backdrop of weak binding sites on binding hotspots. This reduction in the number of parameters greatly simplifies our analysis in the main text.

We assume that there are a small number of strong binding sites, which are bound symmetrically at appreciable rates—termed “hotspots” in the main text—and a larger number of weaker binding sites, which compete with hotspots for PRDM9 binding, but are rarely bound symmetrically. Specifically, we require that weak binding sites are much weaker and more numerous than hotspots, i.e., *k*_2_ ≫ *k*_1_ and *n*_2_ ≫ *n*_1_ respectively, and that the number of weak bind sites far exceeds the number of PRDM9 molecules, i.e., *n*_2_ ≫ *P*_*T*_, such that any given weak binding site is rarely bound, i.e., *H*_2_ ≪ 1. Alongside Eq. S1.6, the latter condition implies that

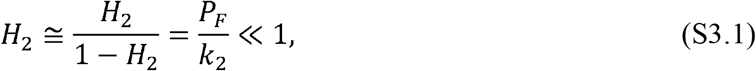

and thus that

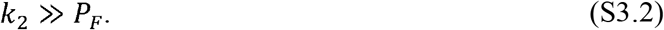

These conditions also imply that the change to the number of weak binding sites is negligible, implying that we can treat *n*_2_ as approximately constant, leaving us with a single variable: the number of hotspots, *n*_1_. We also require that most PRDM9 molecules are bound at equilibrium, i.e., that *P*_*F*_ ≪ *P*_*T*_. In its extreme, this implies that the expected number of free PRM9 molecules in the absence of hotspots 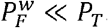.

In order to demonstrate that under these conditions, we can approximate the dynamics in terms of two parameters (P_T_ and 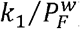), we show that the equations that govern the evolution of a PRDM9 allele and its binding sites are well approximated using these parameters alone. First, with negligible probability of a symmetric DSB at a weak binding site (*α*_2_ ≈ 0), the fitness of a homozygote for a PRDM9 allele is well approximated by

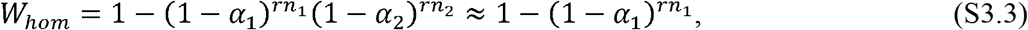

which depends only on n_1_ and on the probability of a symmetric DSB at hotspots, *α*_1_. The same can be easily shown for the fitness of a heterozygotes for PRDM9: namely, that it depends only on *n*_1_ and *α*_1_ of the two PRDM9 alleles. In turn, both the probability of a symmetric DSB at a hotspot (Eq. 4):

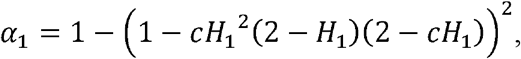

and the rate of gene conversion experienced by hotspots (Eq. 9):

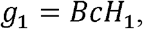

depend on the probability of hotspots being bound, *H*_1_, and on the probability of bound sites experiencing a DSB, *c*. Given that *P*_*F*_ ≪ *P*_*T*_, the probability of bound sites experiencing a DSB is well approximated by

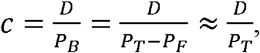

with *P*_*T*_ being the total number of PRDM9 molecules in a homozygotes (the result for heterozygotes for PRDM9 is the same, given our assumption that the sum of the number of PRDM9 molecules of both kinds equals *P*_*T*_; see Eq. 3). Thus, what remains for us to show is that *H*_1_ depends only on *n*_1_, *P*_*T*_ and 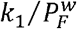.

To this end, we derive an equation for *H*_1_ and show that it can be expressed in terms of these parameters alone. First, we express the total number of PRDM9 molecules as the sum of those that are free, bound to hotspots, or bound to weak binding sites:

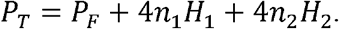

Substituting *H*_2_ ≈ *P*_*F*_ /*k*_2_ (Eq. S3.1) and *P*_*F*_ = *H*_1_*k*_1_/(1 − *H*_1_)(from Eq. S1.6) and rearranging this equation, we find that

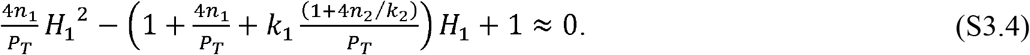

Next, we solve for the number of free PRDM9 in the absence of hotspots. Once again, we express the total number of PRDM9 molecules as a sum of free and bound ones, and substitute *H*_2_ ≈ *P*_*F*_ /*k*_2_ (Eq. S3.1), where in this case, we find that

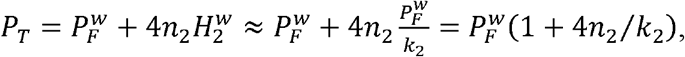

and thus that

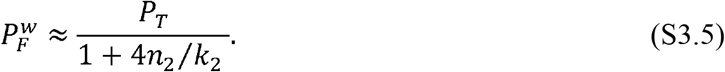

Substitution this expression into Eq. S3.4, we attain a quadratic for *H*_1_,

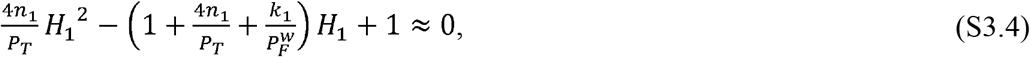

with coefficients that are expressed in terms of, and.

## Supplemental figures

**Figure 3 - Figure Supplement 1:**
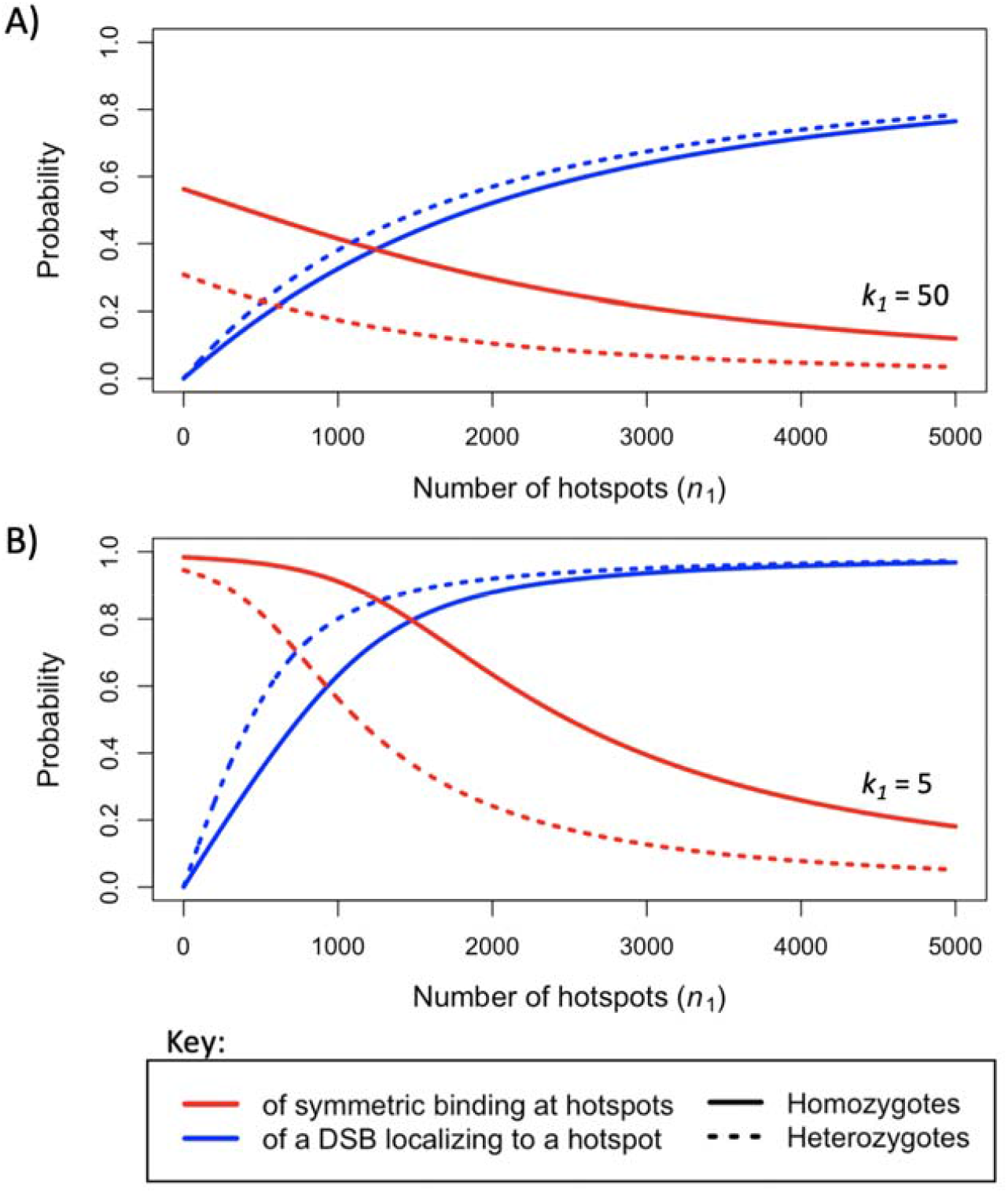
The probability that a hotspot has been symmetrically bound (red), and the probability (or proportion) of DSBs localizing to hotspots (blue), when considering hotspots with (**A**) a relatively strong dissociation constant, or (**B**) a weaker one, under the two-heat model in individuals homozygous (solid lines) or heterozygous (dashed lines) for PRDM9.

**Figure 6 - Figure Supplement 1:**
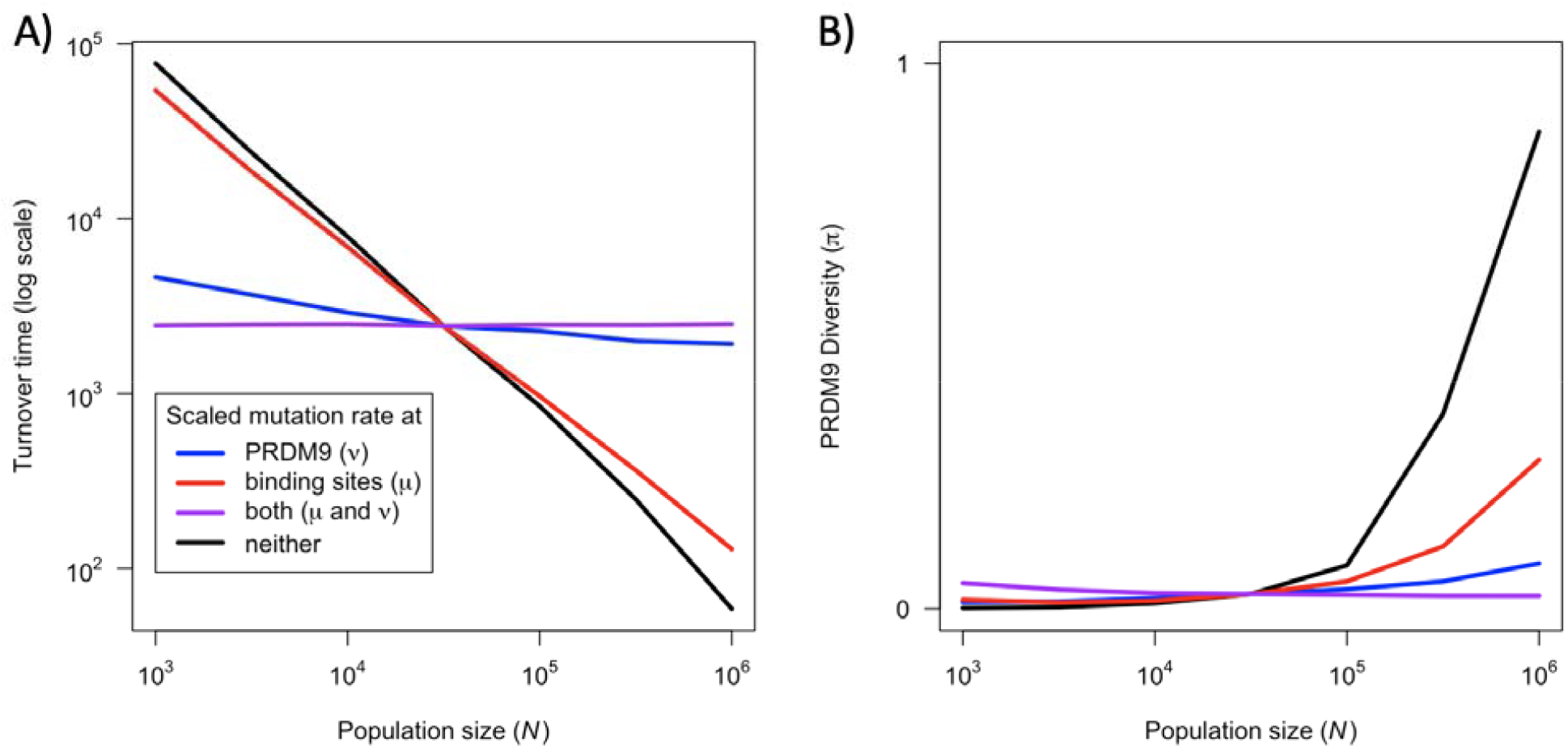
The effect of population size on the rate of turnover (**A**) and diversity (**B**) at PRDM9 while keeping population-scaled mutation rates at PRDM9, binding sites, or both, constant (when).

**Figure 6 - Figure Supplement 2:**
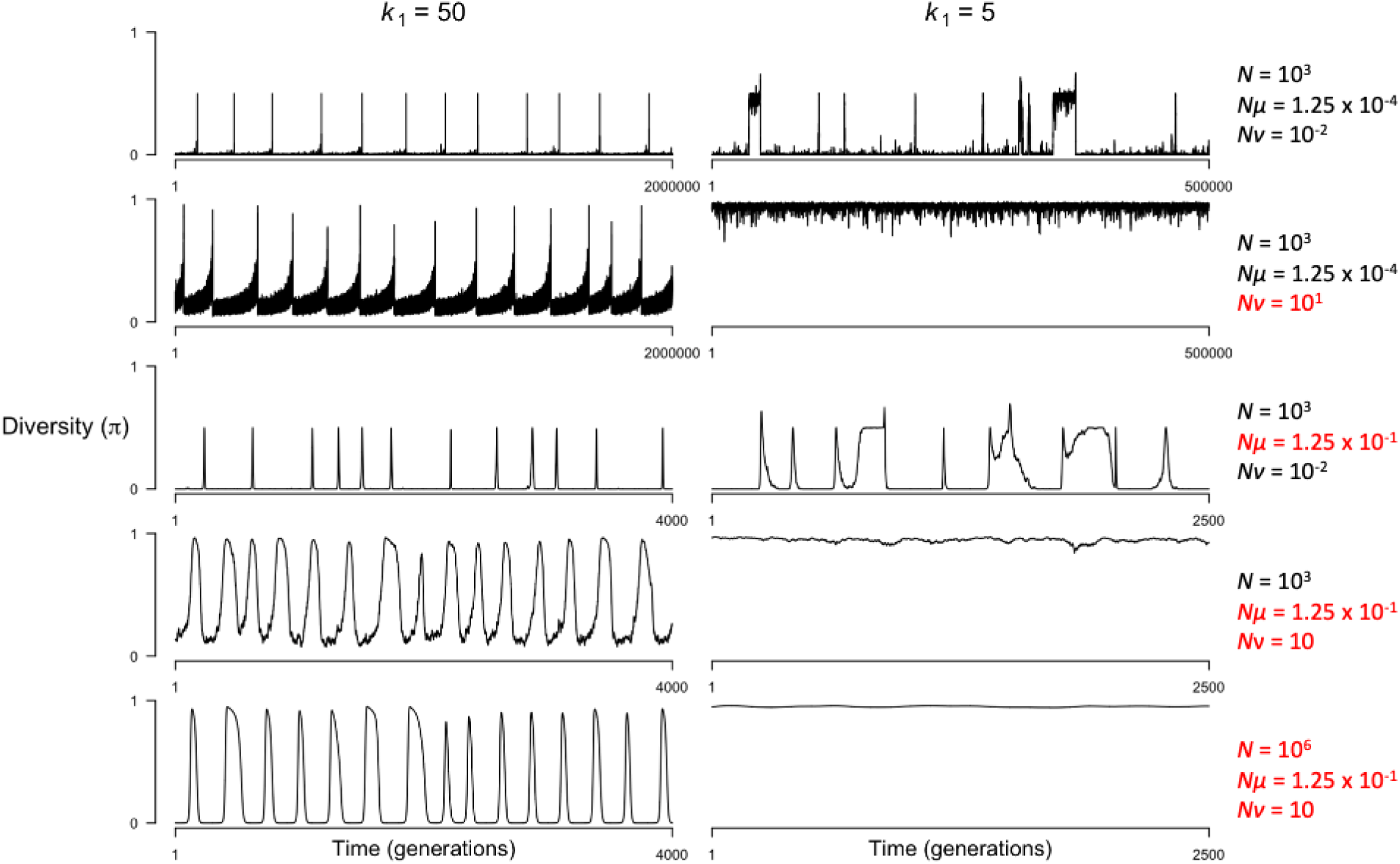
The effect of population-scaled mutation rates on the dynamics of the model when hotspots are cold () or hot (). Values for population size () and mutation rates at PRDM9 binding sites () and at the PRDM9 locus () used in each simulation are shown to the right of each row. Values shown in black indicate those typical for small populations () and values shown in red indicate those typical for large populations ().

## Notes

### Competing Interest Statement

The authors have declared no competing interest.

